# eDNA sampling reveals no negative effects of beaver recolonisation on the catchment-scale distribution of migratory fish

**DOI:** 10.1101/2025.05.13.653709

**Authors:** James A. Macarthur, Alan Law, Nigel Willby, Dasha Svobodova, Nathan P. Griffiths, Roo Campbell, Martin J. Gaywood, Colin W. Bean, Lori Lawson Handley, Melanie Smith, Shaun Leonard, Chris Conroy, Victoria L. Pritchard, Bernd Hänfling

## Abstract

1. Reintroduction of keystone species is considered part of the solution to the current biodiversity crisis. The Eurasian beaver (*Castor fiber*) is one such species, shaping its habitat by felling trees, building dams and creating wetlands. However, while the potential benefits to aquatic biodiversity and ecological functioning have been studied on a local scale, the impacts of beavers on catchment-scale processes such as fish migration remain understudied.
2. Sequencing of environmental DNA (eDNA metabarcoding) from water samples is a cost-effective method to study species distributions across large geographical scales. Here, eDNA samples (*n*=426) were collected from 142 sites across the UK’s oldest and largest established wild beaver population, located on Tayside, East Scotland, and analysed using a vertebrate specific metabarcoding assay. We combined presence/absence data from eDNA results with other environmental and anthropogenic variables to model the effects of beaver presence on the distribution of three migratory fish species.
3. Using generalised linear models, we found no effects of the current beaver presence on the distribution of Atlantic salmon or lamprey, but a positive co-occurrence with European eel at the catchment scale. Model outputs also reinforced previous findings on the impact of barriers to migration and other abiotic and biotic factors on fish species, demonstrating the effectiveness of eDNA sampling in rivers for understanding species distributions at a catchment scale.
4. This case study provides novel insights into the co-distribution of beavers and migratory fish, and highlights how catchment-wide eDNA monitoring can be applied by environmental managers to aid decision-making and impact assessment more generally.

## Introduction

Globally, the average size of monitored wildlife populations in freshwater systems has been estimated to have declined by 85% in the last 50 years (WWF, 2024). The reintroduction of beavers (*Castor spp*.), aquatic mammals once widespread throughout the northern hemisphere, has been proposed as a nature-based solution to address this freshwater biodiversity crisis (Law et al., 2019a; Stringer & Gaywood, 2016). Habitat modifications by beavers, primarily dam building, alter the hydraulic characteristics of rivers and slow the flow of water, thereby converting lotic systems to lotic-lentic mosaics (Naiman et al., 1986). The creation of these heterogeneous wetlands has been shown to benefit biodiversity (Brazier *et* al., 2021; Elmeros *et al*., 2003). However, while the potential benefits to aquatic biodiversity and ecological functioning have been widely promoted at a local scale, the impacts of beavers on catchment-scale processes such as fish migration remain understudied (Brazier et al., 2021; Kemp *et al*., 2012).

Beaver-mediated habitat modifications have the potential to influence migratory fish populations. Negative effects might arise from dams which could restrict or delay the two-way passage of migrating fish, particularly during periods of low flow (Cutting *et al*., 2018; Needham et al., 2025) or reduce the availability of suitable spawning habitat (Kemp *et al*., 2012). Lokteff *et al*., (2013), found interspecific variation in the passability of North American beaver dams by Pacific salmonids in Utah where the movements of native Bonneville cutthroat trout (*Oncorhynchus clarkii utah*) and non-native brook trout (*Salvelinus fontinalis*) were unaffected, although non-native brown trout (*Salmo trutta*) movements were restricted. Similar telemetry studies on bull trout (*Salvelinus confluentus*) in Montana were unable to determine whether dams restricted the upstream distribution of fish or if bull trout actively selected beaver-created wetlands (Marshall Wolf et al., 2024). Research in Norway demonstrated that European beaver dams were unlikely to be a barrier to Atlantic salmon (*Salmo salar*) and brown trout, however salmonid movements were higher in control stretches than beaver dammed reaches (Malison & Halley, 2020). Finally, beaver dam passage for Arctic grayling (*Thymallus arcticus*) in Montana was strongly correlated with hydrological characteristics, temperature and dam status (Cutting et al., 2018). These studies highlight how dam passability varies depending upon stream geomorphology, underlying hydrology, environmental conditions, fish life history, individual fish attributes and the physical characteristics of dams.

Beyond dam construction beaver habitat modifications such as channel excavation, woody debris accumulation, and bank burrowing significantly enhance fish communities including migratory fish by increasing habitat heterogeneity (Kemp et al., 2012). These non-damming activities create pools, backwaters, and undercut banks, offering shelter, spawning grounds, and improved foraging conditions (Pander *et al*., 2025). Such modifications particularly benefit juvenile and small-bodied fish by reducing predation risk and flow stress (Fritz & Gangloff, 2022). Additionally, beaver-driven structural changes promote natural stream processes like sediment sorting and flow variability, which support diverse fish life histories (Bylak et al., 2014; Needham et al., 2025).

The Eurasian beaver (*C. fiber*) has been present in the wild within Tayside, east Scotland since at least 2001 (Campbell et al., 2012). Whilst the exact origin of these beavers is unknown, surveys by the lead Government Agency, NatureScot, estimate that the population has increased from approximately 146 individuals in 2012 to an estimated 954 individuals in 2021 (Campbell-Palmer *et al*., 2021; Campbell *et al*., 2012). Meanwhile, the number of adult Atlantic salmon returning to Scottish rivers have continued to decline, which has prompted the IUCN to reclassify the British salmon population from Least Concern to Endangered (Darwall & Noble, 2023). In 2015, a mapping study estimated there would be a 72% habitat overlap between beavers and salmon across the Tay catchment (Beaver Salmonid Working Group, 2015), and there remains uncertainty surrounding the future of these two species of conservation concern and their interactions.

Whilst beaver and salmon interactions remain understudied in Scotland Needham *et al*., (2021) found that beaver modifications to a Scottish stream benefited the local brown trout populations where the beaver-modified habitats supported higher abundances of larger trout and a wider variety of age classes. Subsequent telemetry studies highlighted that brown trout could successfully navigate the beaver dams, although successful passage was determined by environmental conditions and individual fish characteristics (Needham et al., 2025). These studies were, however, limited to a small number of streams and focused primarily on potamodromous brown trout. Meanwhile the effects on conservation priority migratory species such as brook lamprey (*Lampetra planeri*), European river lamprey (*Lampetra fluviatilis*) and critically endangered European eel (*Anguilla anguilla*), which are particularly vulnerable to changes in water quality and habitat connectivity, remain unknown (Nunn *et al*., 2023). Globally, beaver distributions are expanding through conservation translocations (Elmeros *et* al., 2003), legal protection and natural colonisation (Campbell-Palmer et al., 2021). Therefore, the need to understand the impact of expanding beaver populations on migratory fish species is becoming increasingly important.

Environmental DNA (eDNA) has become an important monitoring tool to assess the status of freshwater biodiversity (Hänfling et al., 2016; Pont et al., 2018). The development of high-throughput DNA sequencing of target regions (“metabarcoding”) provides a cost-effective method to study species distributions across a large-scale (Broadhurst *et al*., 2021; Griffiths et al., 2023). eDNA has also been shown to increase detection of rare fish species such as European eel when compared with traditional fish survey techniques (Griffiths *et al*., 2020). However, whilst eDNA has often been applied at a local scale within lakes (Hänfling et al., 2016), ponds (Harper *et al*., 2018) and rivers (Broadhurst et al., 2021; Pont et al., 2018) its potential for upscaling across entire catchments is only just starting to be realised (Blackman et al., 2024).

In the current study we applied a catchment-scale eDNA survey across the Tayside region to characterise the co-distribution of beavers and migratory fish species. Specifically, we modelled the occurrence of beaver eDNA, alongside key anthropogenic (artificial migration barriers), environmental (habitat characteristics) and biotic (predator presence) covariates on the occurrence of salmonids, lamprey and eel, across Tayside, to investigate if the presence of beavers affects the distribution of migratory fish at the catchment scale.

## Methods

### Site selection

The Tayside region was gridded to ensure balanced spatial coverage, before 142 sites near beaver activity were selected (Fig. 1) based upon a catchment-wide field signs survey carried out in 2021 (Campbell-Palmer *et al*., 2021). In most river systems, three sites were selected upstream of beaver activity at 5 km intervals depending on the nearest road access. To identify river systems where beaver populations were likely to have expanded since the 2021 survey, we integrated the data layer of ‘Potential Core Beaver Woodland’ produced by NatureScot to prioritise sites which had suitable beaver habitat (Stringer et al., 2018).

**Figure 1:**
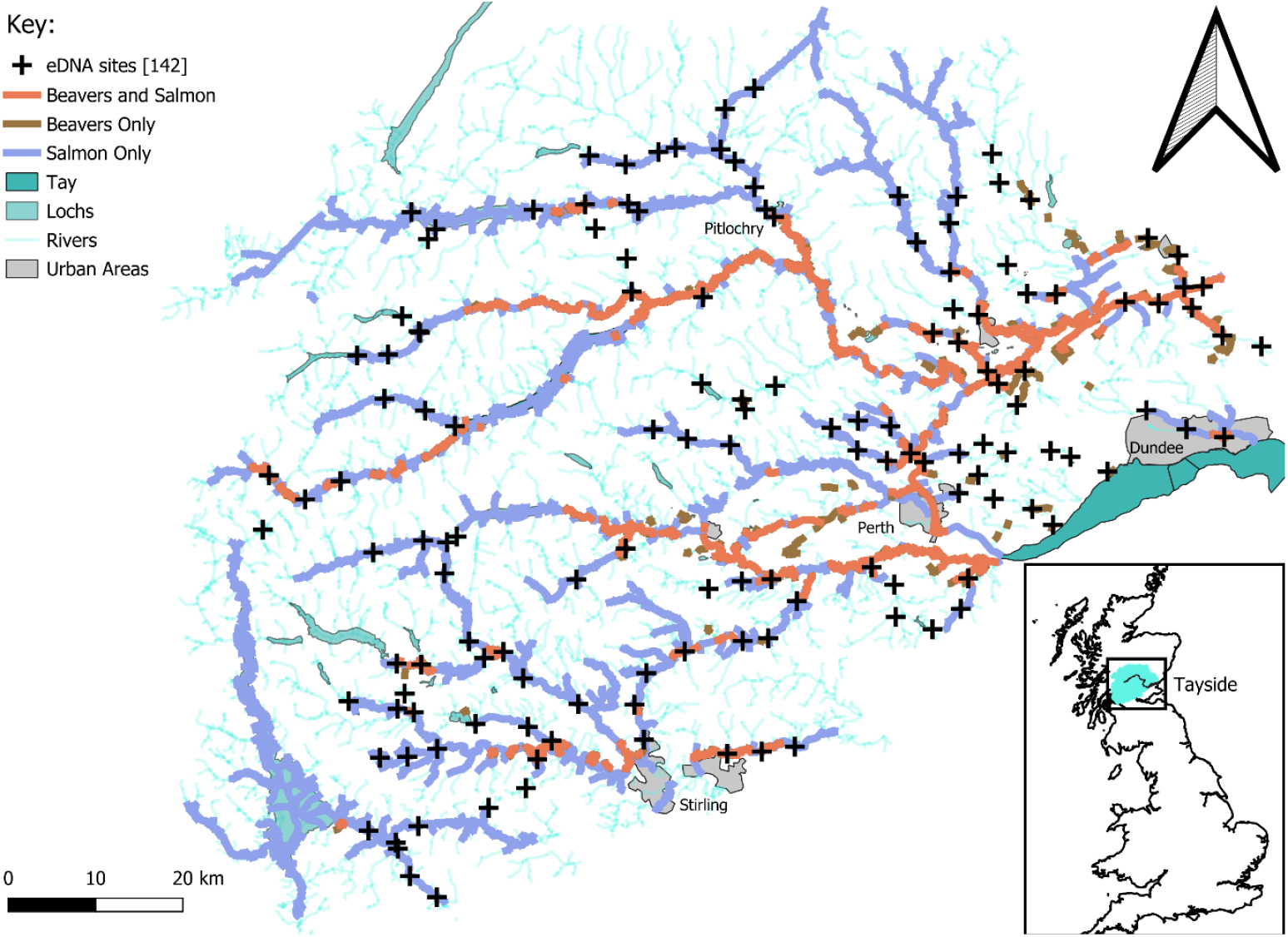
eDNA sites sampled across Tayside (n =142; black cross). The beaver distribution is taken from the 2021 NatureScot survey (Campbell-Palmer *et al*., 2021). The salmon distribution is informed by the “Salmon rivers in Scotland” (Marine Directorate, 2008), which is based on digital spatial data licensed from the UK Centre for Ecology & Hydrology, © NERC (UKCEH) and contains Ordnance Survey data © Crown copyright and database rights 2024.

The “Salmon Rivers in Scotland” layer (Marine Directorate, 2008) which has collated known information on the presence of salmon since the 1980s and is an update of (Gardiner & Egglishaw, 1986) using the UKCEH digital river network (Moore et al., 1994), was used to select sites that overlapped with beavers. Electrofishing data from the NEPS (National Electrofishing Programme for Scotland) surveys were used to provide quantitative information on juvenile salmonid densities that could be used to prioritise sampling productive salmon habitats (Malcolm et al., 2023). Finally, the distribution of other priority migratory fish species were taken from the 2018 NEPS electrofishing fish counts (Malcolm et al., 2019b). Of the 53 electrofishing records which overlapped the current study, eels and lamprey were detected in 11 and 4 sites, respectively.

### Sample collection and filtration

Between 06/07/2023 – 18/08/2023, 426 surface water samples were collected at 142 sites, following the methodology described in Griffiths *et al*., (2023). Field blanks (n = 19) of purified water were used daily to monitor contamination. All water samples were vacuum-filtered within 24 hours of collection through sterile 0.45 μm mixed cellulose nitrate membrane filters with pads (47 mm diameter; Whatman, GE Healthcare) using Pall filtration units and then stored at -20 °C. See supplementary-1 for full details. Ethical approval was obtained under the application “ETH2223-0838” to UHI.

### DNA extractions

DNA was extracted using the mu-DNA lysis water (Sellers et al., 2018) with modifications: during the lysis stage, instead of using garnet beads, 34 μl of 20 mg/mL proteinase K (VWR, Leicestershire, UK) was added to 750 μl of lysis solution and 250 μl of water lysis additive and incubated at 55°C overnight. Extraction blanks (n = 20) were used to monitor contamination.

### Metabarcoding

Library preparation followed a modified Griffiths *et al*., (2023) protocol. Nested metabarcoding followed a two-step PCR approach in which both PCRs used multiplex identification tags to enable sample identification (Kitson et al., 2019). A positive control (quantified at 0.05 ng/μl) from the cichlid (*Astatotilapia calliptera*) that is not found in the UK and a negative control of molecular grade water were used for each sub-library. The first PCR was carried out in triplicate (3x PCR replicates) to amplify a 106bp fragment using published 12S ribosomal RNA primers (Kelly et al., 2014; Riaz et al., 2011). These primers have been validated both *in vitro, in silico* and *in situ* for UK vertebrates (Hänfling et al., 2016), confirming that for the purpose of the current study, all UK migratory fish species can be detected. However, they cannot distinguish between river or brook lamprey which are therefore assigned to the genus *Lampetra*.

Sub-libraries were cleaned using the ProNex® Size-Selective purification beads at a ratio of 1.5x to 50 μl. A second PCR was carried out in duplicate to bind pre adapters, indexes, and Illumina adapters to each sub-library before a double size selection clean-up (Quail et al., 2009) was carried out at the ratios of 1.2x and 0.4x magnetic beads to 50 μl of PCR product.

Cleaned sub-libraries were quantified and diluted to 5 ng/μl before they were pooled proportionally based on the number of samples. The pooled libraries were then diluted to 4 nM and quantified via qPCR using the Collibri™ Library Quantification Kit for Illumina (Roche, Hertfordshire, UK). The final libraries were denatured and sequenced at 13pM with 10% PhiX Control on an Illumina MiSeq using a MiSeq Reagent Kit v3 (600 cycle) (Illumina Inc., San Diego, CA, USA).

### Data filtering

Reads were demultiplexed using a custom script then processed with Tapirs, (https://github.com/EvoHull/Tapirs), for quality-trimming, merging and taxonomic assignment using a curated UK vertebrate reference database (Harper et al., 2018). Taxonomic assignment followed a lowest common ancestor approach based on basic local alignment search tool (BLAST) matches with minimum identity set at 98% (Griffiths et al., 2023).

Following the bioinformatics, a minimum threshold of five reads was applied to remove low-frequency reads alongside a species-specific contaminant threshold. This was applied on a per flow cell basis to remove any reads in samples which were lower than detections in controls. Post-filtering, field replicates were combined, into site-level presence (1) or absence (0).

### Environmental metadata and covariates

Sixteen covariates that could explain the distribution of the three migratory species were selected from literature, based on available GIS data (Ferreira et al., 2013; Griffiths et al., 2023; Malcolm et al., 2019a). These included migration barriers (SEPA-WMS, 2018), geospatial variables related to the site’s location (Davies et al., 2022; Eurogeographics, 2023), adjacent land use (Morton et al., 2024), physico-chemical parameters (Hollis *et al*., 2019; Johnson et al., 2005), and the presence of other key species such as predators (based on this studies’ eDNA data). See supplementary-2 for full details. Artificial and natural barriers downstream were combined before being reclassified into three types following SEPA’s guidelines: “passable barriers with fish passes”, “passable barriers without fish passes” and “impassable barriers” (SEPA-WMS, 2018).

Land Cover Maps (Morton et al., 2024) were used to calculate upstream land-use within a 25 m buffer either side of all contiguous rivers located within a buffer extending 1 km upstream of each site. To simplify the land-use categories several composite groupings were created Urban (Urban + Suburban), Wetland (Fen, Marsh and Swamp + Bog), Acid Moorland (Acid Grassland + Heather Grassland + Heather and Shrub), Agricultural (Arable + Improved Grassland). Freshwater land-use was omitted from this analysis to minimise noise as the rivers sampled were often too small to be recorded on the land cover map.

The presence of Northern pike (*Esox lucius*) was derived from the 2023 eDNA samples to assess the effect of the overlap of a potential predator species on salmon. Other predator species such as Eurasian otter (*Lutra lutra*) were not included in this analysis as they have a low detection probability based on eDNA sampling (Broadhurst et al., 2021). Lastly, the distribution of beavers was incorporated from the 2023 eDNA surveys as 1 (presence) and 0 (absence) to investigate the effects of beaver presence on the distribution of each migratory fish species.

### Model selection

PCA was used to reduce land-use, watershed and climate covariate complexity. Correlation matrices were used throughout to ensure highly correlated variables were omitted and a VIF (variance inflation factor) threshold of < 5 was used to avoid multicollinearity in the final models. Channel slope and mean atmospheric temperature, bankfull river width and acid moorland were all omitted from the final models as they were highly correlated with elevation, upstream catchment area and impassable barriers respectively. All continuous variables except for pH and distance from coast were log transformed to improve their distribution.

Variables were mean centred to 1 SD and scaled to improve comparability and reduce the impacts of outliers (Law et al., 2019b).

A series of binomial generalised linear models (GLMs) were used to investigate the effects of covariates on the presence of migratory fish species. A reverse stepwise approach was used to select the most parsimonious model for each species based on the lowest Akaike Information Criterion (AIC). Initially, catchment identity was added as a random effect in GLMMs using the package “lme4” (Bates et al., 2015), however it did not improve the model AIC and was therefore removed. We considered all variables which were retained following the model selection to be meaningful, but also report statistical significance at the 95% confidence level to highlight where relationships exhibit strong evidence versus weaker evidence (retention in the most parsimonious model but with P>0.05).

The full models were:

**Migratory species ∼ Impassable barriers + Passable barriers with fish passes + Passable barriers without fish passes + Upstream catchment area + Total rainfall + Elevation + pH + Distance from coast + Beavers + Pike + Agricultural + Broadleaved Woodland + Coniferous woodland + Urban + Wetland**

All GIS and covariate extraction was done in QGIS (QGIS Development Team, 2025), analyses and graphical visualisation used R v.4.5.0 (R Core Team, 2025) with ggplot2 (Wickham, 2016) and sjPlot (Lüdecke, 2024).

## Results

Beavers were detected in 72 (50.7%) sites, of which 79% were positive across all three replicates and 28/34 (82.4%) sites with beaver field signs from 2021 within a 500 m circular buffer were positive. Four migratory fish species: brown trout, salmon, eels and lamprey (*Lampetra spp*.) were detected across the 142 sites. Sea lamprey (*Petromyzon marinus*) were not detected in the current study. There was considerable overlap between beavers and migratory fish across Tayside (Table 1, Supplementary 11-14). Brown trout were detected at 138 (97.2%) sites which meant there were insufficient absences to model their distribution. Salmon were detected in 93 (65.5%) sites, of which 88% were positive across three replicates (Supplementary 10). Lamprey were found at 69 (48.6%) sites (84% positive across three replicates) and eels at 103 (72.5%) sites (87% positive across three replicates). Occurrence of these three species was therefore modelled (Fig. 2, Supplementary-3-9)

**Table 1:**
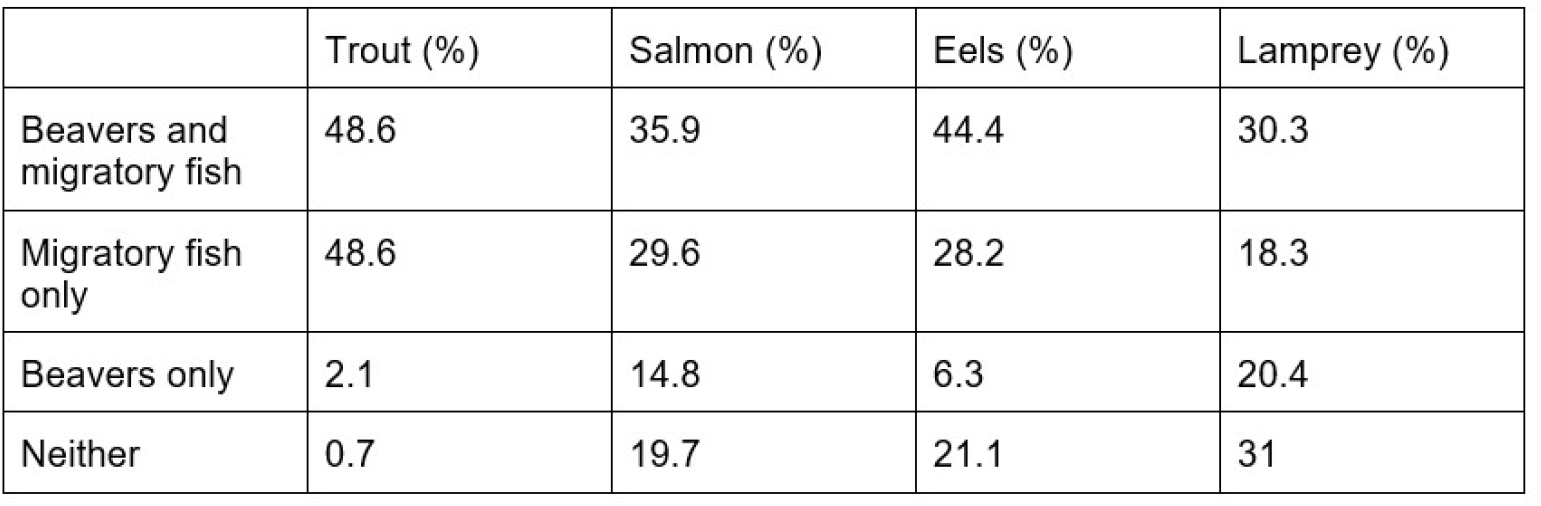
Percentage overlap between beavers and migratory fish across the 142 eDNA sites

**Figure 2:**
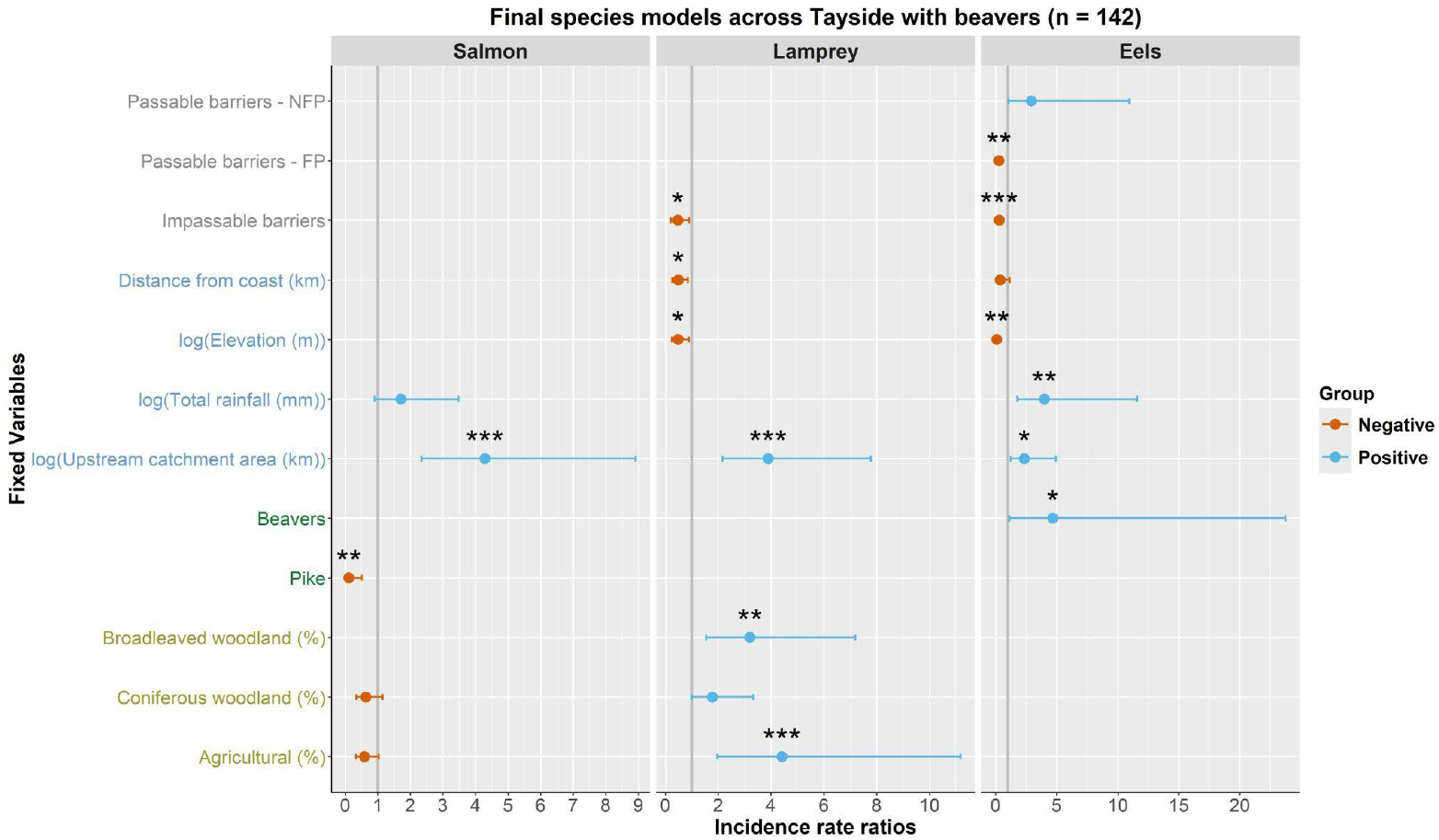
Final model outputs for the migratory fish species. Lines are coloured by effect (Negative = Orange, Positive = Blue) and asterisks denote the level of significance (* =0.05, ** =0.01, ***=0.001). Covariate text is coloured by type (Barriers = Grey, Physical = Blue, Species = Green, Land-Use = Yellow) and the whiskers represent the 95% confidence interval.

There was no evidence for an effect of beaver presence on the distribution of salmon or lamprey, with beavers not included in the final model for both species. However, a positive co-occurrence was demonstrated between beavers and eels (P <0.05) (Supplementary-6). Contrastingly, the presence of pike had a moderately negative effect on the distribution of salmon (P <0.01).

Impassable barriers had a consistent negative effect on all three migratory fish species (eels: P <0.001; lamprey: P <0.05). No salmon were detected upstream of impassable barriers, therefore the 22 sites upstream of impassable barriers were omitted from the salmon models. Barriers with fish passes also had a strong negative effect on the distribution of eels (P <0.01). The distribution of all three migratory fish species was positively correlated with upstream catchment area (salmon: P <0.001; lamprey: P <0.001; eels: P <0.05). Lamprey and eel presence were both negatively correlated with elevation (lamprey: P <0.05; eels: P <0.01). Distance from the coast had a negative effect on the distribution of lamprey (P <0.05) and eels had a positive association with total rainfall (eels: P <0.01).

The percentage cover of agricultural land had the largest positive effect of land-use on lamprey distribution (P <0.001). Broadleaved woodland (P <0.01) was also positively associated with the presence of lamprey. Land-use had no effects on eel presence and was dropped from the final model.

Finally, there was some evidence through their retention by AIC, that the following variables had an effect on migratory fish species distributions, however these effects were non-significant at the 95% level in the final models (Fig. 2): For eels, passable barriers without fish passes (positive, P =0.073) and distance from the coast (negative, P =0.111). For salmon total rainfall (positive, P =0.113), coniferous woodland (negative, P =0.137) and agricultural (negative, P =0.071). For lamprey, coniferous woodland (positive, P =0.057).

## Discussion

This study provides a novel, catchment-scale investigation into the effects of beaver presence on migratory fish distribution using eDNA metabarcoding. By integrating species presence data with environmental and anthropogenic covariates, our results offer new insights into the coexistence of beavers and fish species, particularly in the context of conservation translocation initiatives. Alongside this, these models revealed effects of both natural and anthropogenic barriers, distinct environmental and land-use correlates, and predator interactions consistent with existing literature exemplifying the effectiveness of eDNA sampling in rivers as a basis for understanding species distributions and their drivers.

### Interactions between beavers and migratory fish

We found no evidence that beaver presence negatively affects the distribution of salmon, trout, eels or lamprey across Tayside, despite the ongoing expansion of beaver populations (Campbell-Palmer et al., 2021). A number of prior studies corroborate these findings and demonstrated that the distribution and movement of trout in Scotland (Needham et al., 2021, 2025) and juvenile salmonids in Norway (Malison & Halley, 2020) were not negatively affected by beaver dams. However, these studies were limited to a small number of streams or particular sets of dams, constraints that make it difficult to infer landscape-scale impacts.

Beaver impacts are highly variable influenced by the surrounding environmental conditions, underlying hydrology, beaver densities and habitat characteristics, all of which determine the type, size, stability, and longevity of their habitat modifications (Cutting et al., 2018; Lokteff *et* al., 2013; Marshall Wolf et al., 2024; Naiman et al., 1986). In contrast, our study investigated the impact of beavers on catchment-scale processes such as fish migration where the current co-existence of both beavers and migratory species is evident. It is important to note that few beaver dams were observed in the current study as is typical of large or higher gradient riverine systems. Dams are also regularly removed as part of conflict mitigation (Campbell-Palmer *et* al., 2021), therefore the full effects of beaver activity on migratory fish in Tayside are unlikely to currently be captured and might change as the beaver population expands.

Past studies on migratory fish species have primarily focused on salmonids (Cutting et al., 2018; Lokteff et al., 2013; Malison & Halley, 2020; Needham et al., 2021, 2025), whilst eDNA enabled us to obtain an unbiased and accurate distribution of non-salmonid fishes such as lamprey and eel. The finding that beavers are positively co-distributed with eels across Tayside is consistent with research from Elmeros *et al*., (2003) who suggested that due to their leaky nature, beaver dams were not barriers to eels. In fact, it is likely that the modifications by beavers will provide increased habitat complexity through higher in-channel coarse wood and connectivity to the floodplain which would benefit eels (Harwood et al., 2022).

### Other drivers of fish distribution

The consistent negative effect of impassable barriers on migratory fish species demonstrates the utility of eDNA in confirming the passability of both natural and artificial barriers downstream. The degree to which impassable barriers affected each species varied; whilst salmon could not navigate any of the 22 barriers, lamprey and eels were detected upstream of two and six barriers respectively. This showcases the importance of classifying barrier passability relative to individual species. The complete absence of salmon eDNA above these barriers is unsurprising, as the barrier classification was largely based on knowledge of existing salmon distribution (Gardiner & Egglishaw, 1986; SEPA-WMS, 2018). The presence of lamprey eDNA upstream of two waterfalls likely reflects freshwater-resident populations of brook lamprey. However, these waterfalls are probably impassable to the anadromous river lamprey as recorded on the Endrick Water at the Pots of Gartness (Bull *et al*., 2016). Unfortunately, our current eDNA assay cannot distinguish these two closely related species.

Notably, our results indicated a negative effect of barriers with fish passes on the distribution of eels. This aligns with previous research showing that hydropower infrastructure can hinder the upstream migration of juvenile eels (elvers) (Feunteun, 2002). Whilst elver passes can improve access upstream (Solomon & Beach, 2004), the broader challenge is to ensure suitable downstream migration opportunities for adult silver eels (Piper et al., 2013). The current approach in Scotland is that since safe downstream passage through hydropower dams cannot be enabled, it is better to restrict access upstream entirely. However, with the continued decline of European eels throughout their range (Nunn et al., 2023), and the detection of eels upstream of these barriers in our present study, downstream passage solutions should be considered. This reinforces the need for species-specific passage solutions that account for the unique migratory behaviours and vulnerabilities of different life stages.

Our findings reveal distinct environmental and land-use correlates for species distributions, emphasising the need for species-specific conservation strategies. The positive association between upstream catchment area and the presence of all three species, salmon, lamprey, and eels, highlights the importance of larger, well-connected river systems in supporting migratory pathways and habitat availability. Distance from coast and elevation were both negatively associated with eel and lamprey presence. This is consistent with Griffiths *et al*., (2023) who found that eel presence in Cyprus declined with distance from coast, number of barriers and elevation. However, for lamprey, research from Portugal suggests that this relationship is a product of the shallow gradient microhabitats created in these conditions which provide a favourable silty substrate for ammocoete development and feeding (Ferreira et al., 2013). The distribution of lamprey across Tayside was positively correlated with agricultural, broadleaved and coniferous woodland cover which could all benefit ammocoetes through increased fine sediment (Bull et al., 2016). In contrast, there was some evidence that agricultural land had a weak negative effect on salmon presence, likely because fine sediment deposition could render spawning gravels unsuitable for salmonids (Soulsby et al., 2001). Meanwhile eel distributions were not influenced by land-use, perhaps reflecting their high level of plasticity to environmental variation (Enbody et al., 2021).

Excluding sites upstream of impassable barriers; salmon were absent from six of the 19 sites across Tayside in which pike were detected. This negative effect could be a direct result of predation as pike are known to predate migrating salmonid smolts (Falkegård *et al*., 2023) or it could reflect differing habitat requirements of the two species as pike typically inhabit slower flowing systems (Beaver Salmonid Working Group, 2015) whilst salmon utilise faster flowing rivers (Soulsby *et al*., 2001). Beaver ponds on smaller water courses could support more predatory fish such as pike or piscivorous brown trout; therefore predation of juvenile Atlantic salmon might become more of an issue as the Tayside beaver population expands (Beaver Salmonid Working Group, 2015).

### Limitations of eDNA

The application of eDNA metabarcoding at the catchment scale represents a significant methodological advancement in ecological monitoring. However, several limitations must be acknowledged.

First, our study infers only the presence or absence of species, rather than providing semi-quantitative estimates of abundance. While a clear correlation between DNA sequence abundance and relative fish abundance has been demonstrated (Di Muri et al., 2020; Griffiths et al., 2020), this relationship has not yet been established for other vertebrate species. As such, our conclusions are limited to the spatial distribution of species across the catchment, and we are unable to assess the effects of beavers or other environmental variables on the abundance of migratory species. Additionally, eDNA does not allow for the identification of life stages, therefore the detection of a species is treated as a composite signal encompassing all life stages. Consequently, we are unable to differentiate life-stage-specific habitat preferences (Harwood et al., 2022). We recommend that eDNA monitoring be complemented with targeted electrofishing surveys to assess changes in fish population dynamics and resilience following beaver-related habitat modifications.

Finally, the downstream transportation of eDNA may lead to false positives. In lotic systems, DNA can be carried from its source for distances ranging from hundreds of meters in small streams to over 100 km in very large rivers (Pont, 2024; Pont et al., 2018). While transport distances vary with river size and flow conditions, the rivers in our study are small to medium-sized, disrupted by artificial barriers, and were sampled during low-flow summer conditions, which reduces the likelihood of extensive eDNA transport. Although we cannot entirely rule out occasional long-distance transport beyond our 5 km sampling intervals, we therefore consider this unlikely to be a major source of error. Regarding beaver–migratory fish interactions, the detection of migratory fish upstream of beaver activity suggests these species had successfully traversed beaver-occupied areas downstream to reach the sampling site.

### Wider applications

This study highlights the value of catchment-scale eDNA metabarcoding for assessing species distributions and interactions, offering a practical tool to evaluate the ecological impacts of beaver reintroductions and other environmental factors on migratory fish. Traditional methods for studying barrier effects on fish migration, such as telemetry, electrofishing, or observation, are often costly, spatially limited, and potentially disruptive (Kemp & O’hanley, 2010). eDNA provides a non-invasive alternative, where discontinuity in species detections upstream versus downstream of a structure may signal an unrecognized migration barrier (Laramie *et al*., 2015). With its capacity for repeated, high-resolution sampling, eDNA also enables tracking of seasonal migration and evaluation of barrier mitigation efforts.

Our findings inform management strategies as beaver populations grow across Europe. The lack of negative effects on salmon, brown trout, and lamprey suggests beaver reintroductions can align with their conservation, especially where habitat connectivity is maintained. The positive link with eels also points to potential habitat benefits for declining species. From a policy standpoint, these results support continued beaver translocation to appropriate sites, with attention to local ecological dynamics. Enhancing fish passage infrastructure remains essential to maximize the ecological gains of beaver reintroductions while safeguarding migratory fish.

## Supporting information

Word document containing all supporting information for the publication

## DATA AVAILABILITY STATEMENT

All scripts and corresponding data have been archived and made available at Zenodo:

https://doi.org/10.5281/zenodo.15395641

---

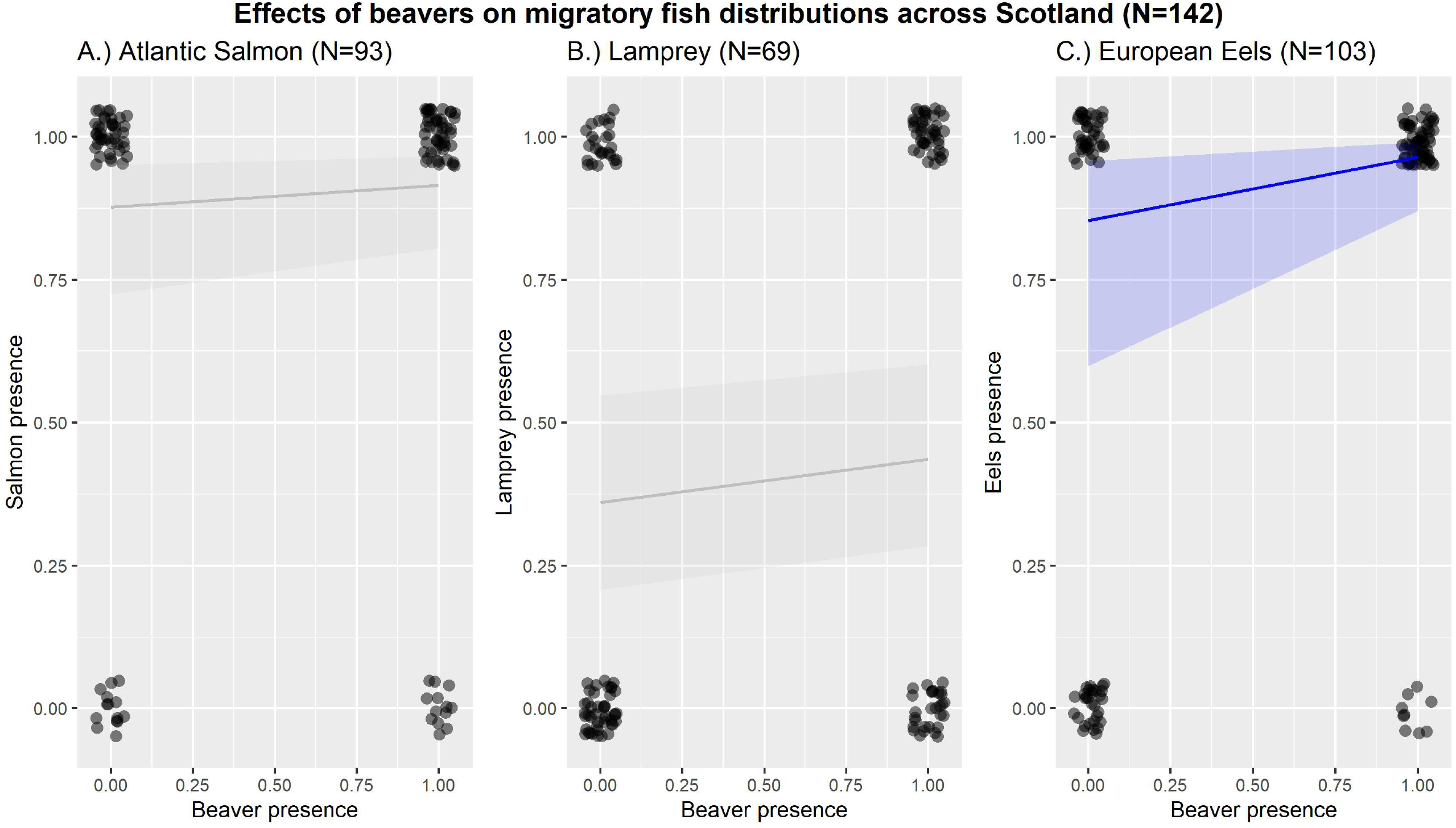

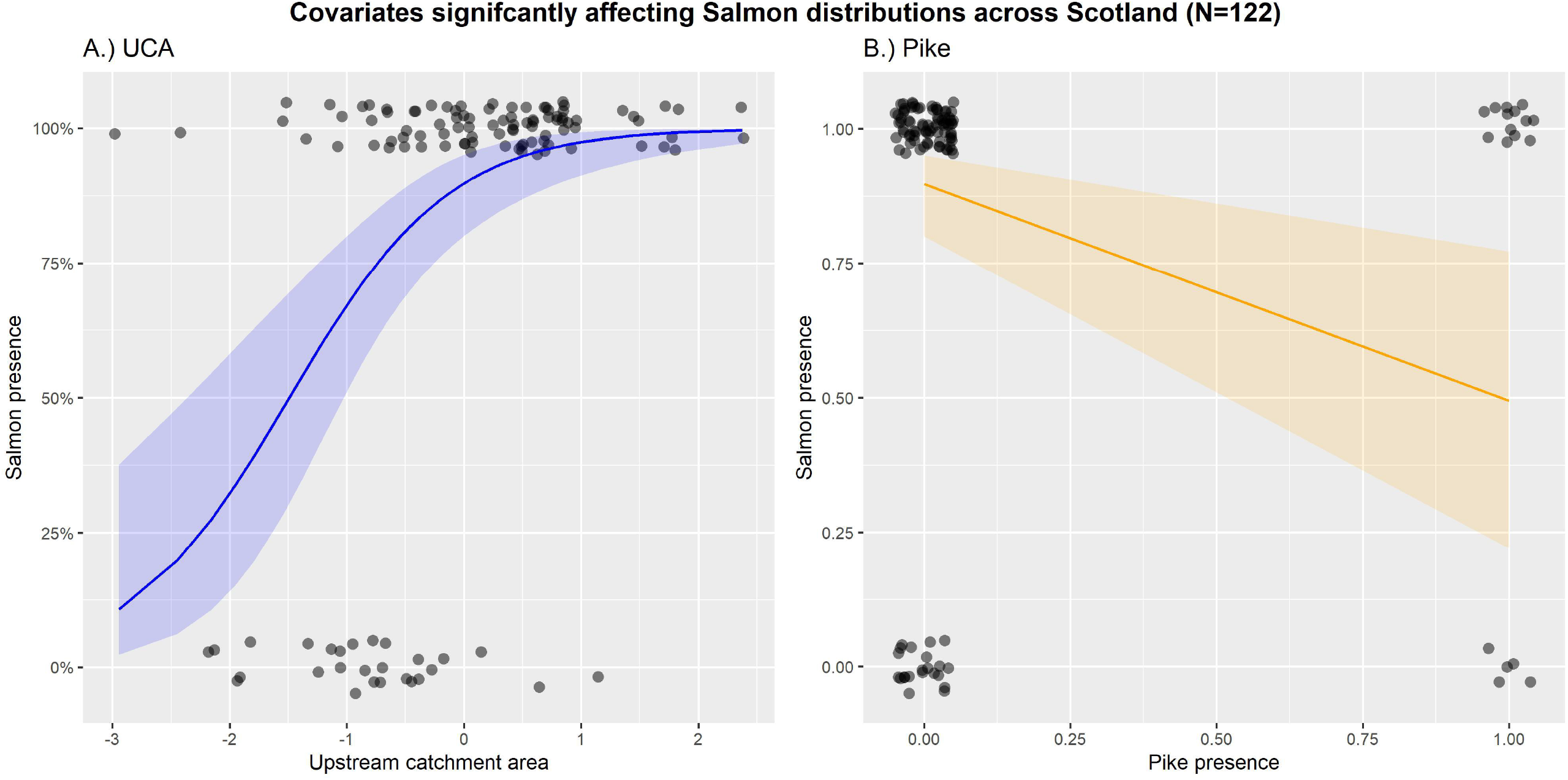

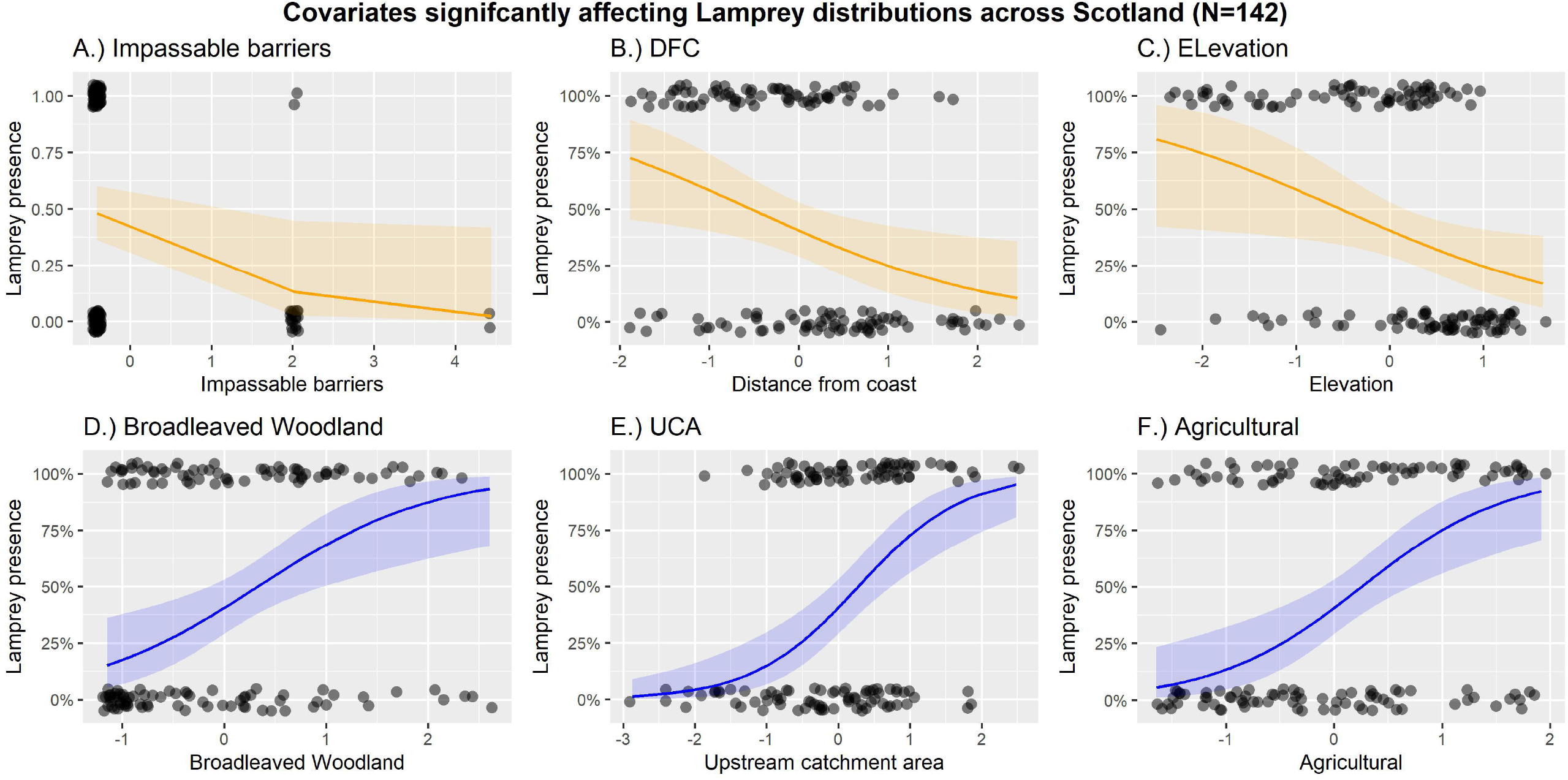

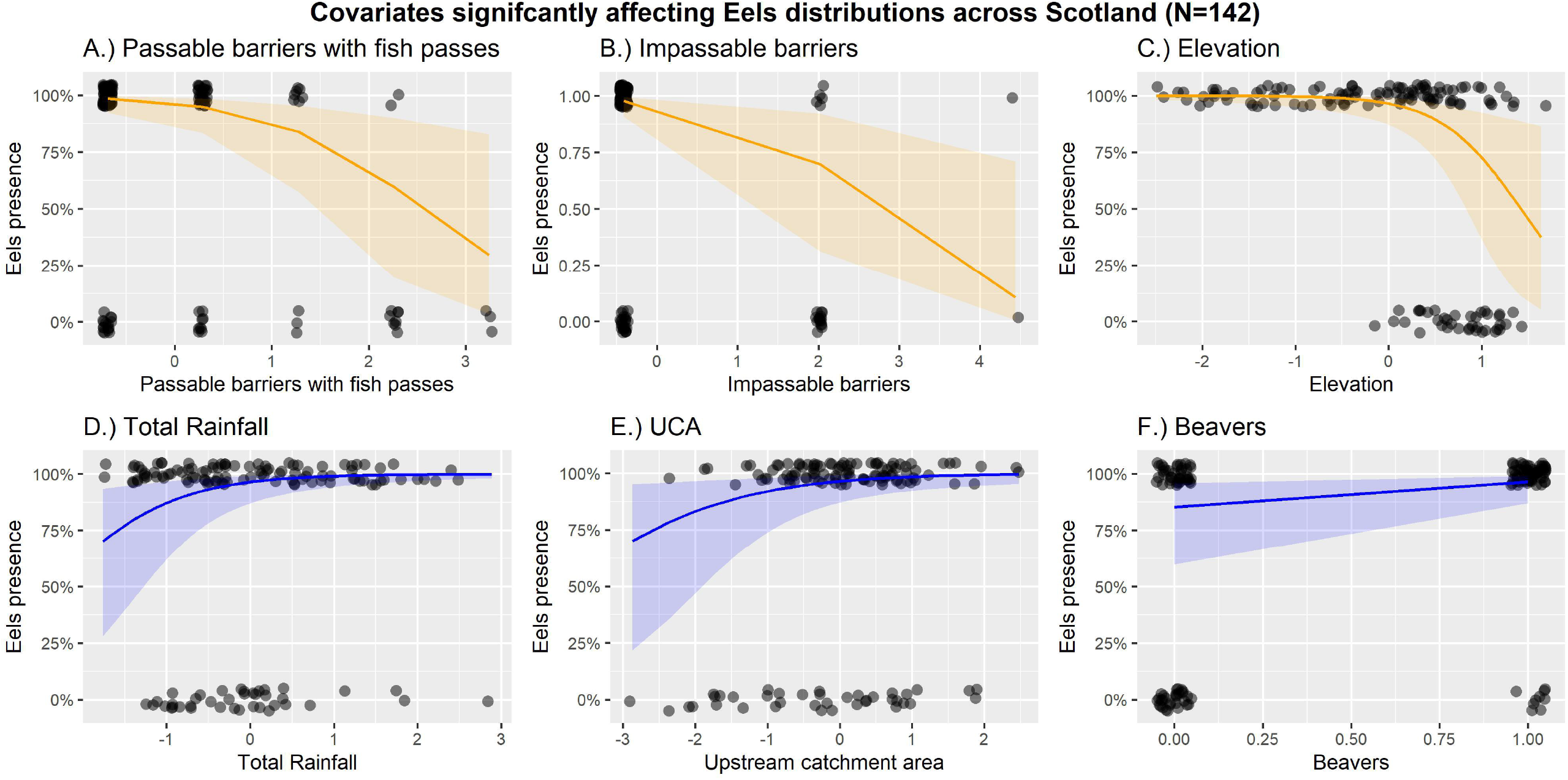

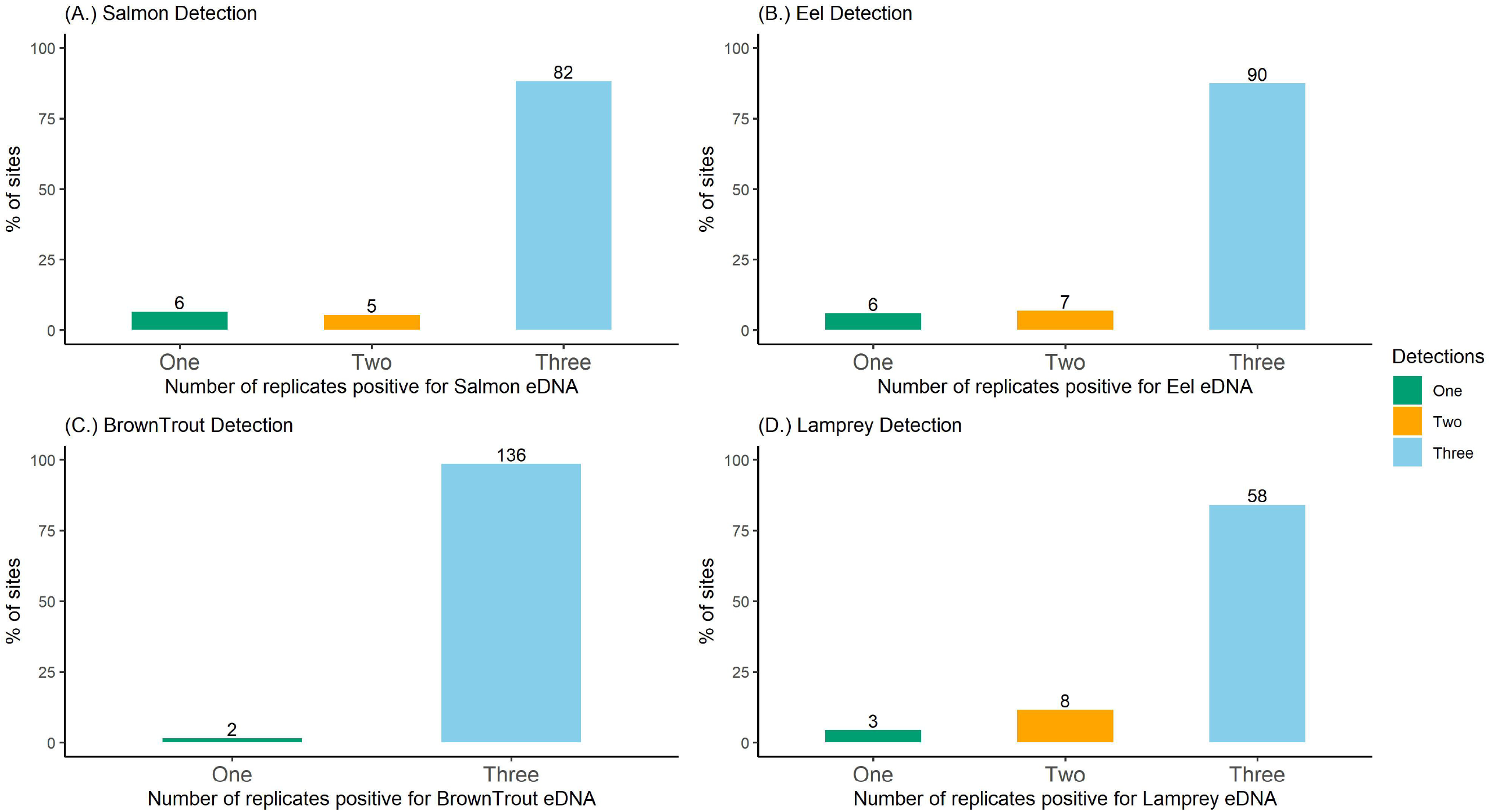

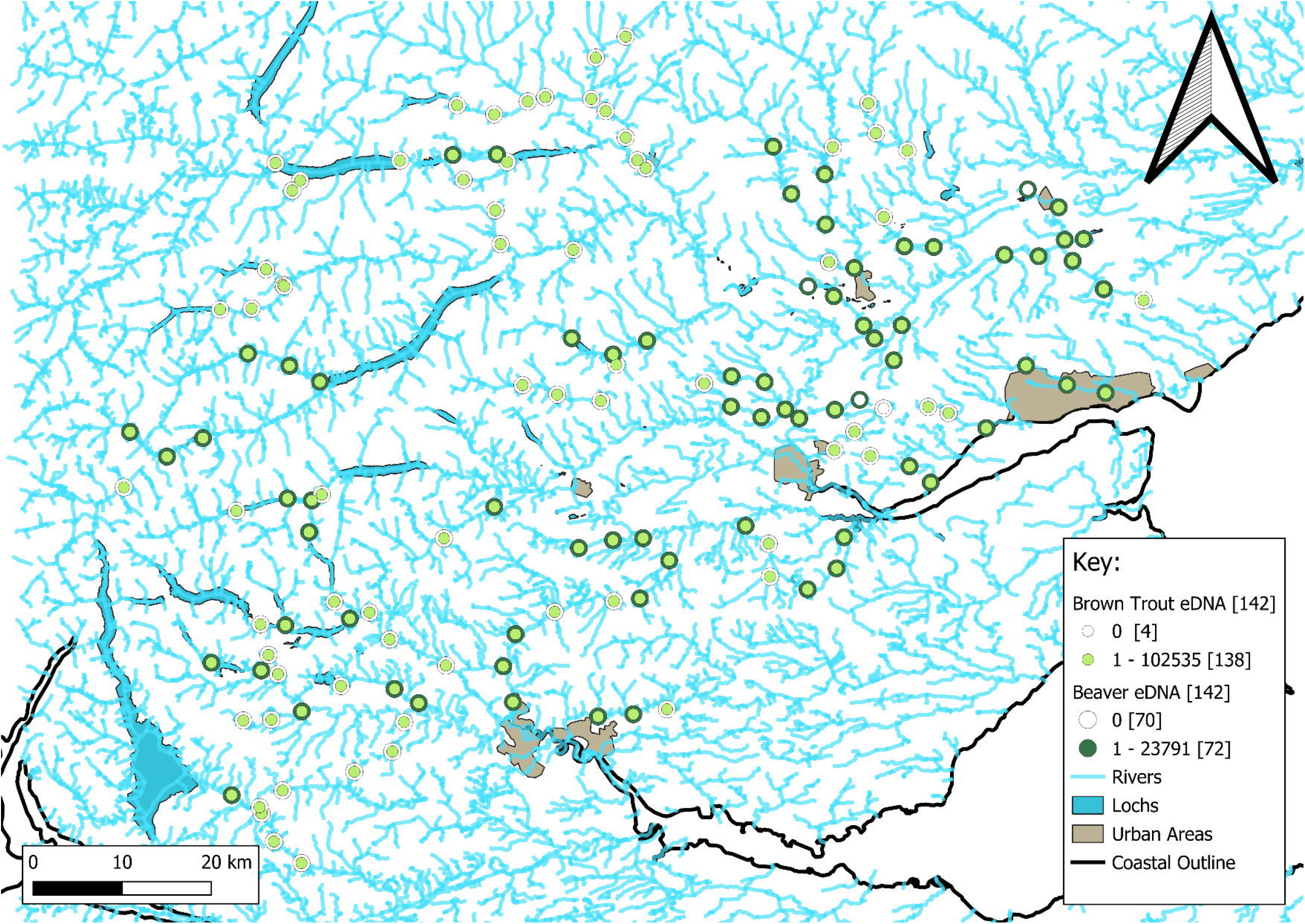

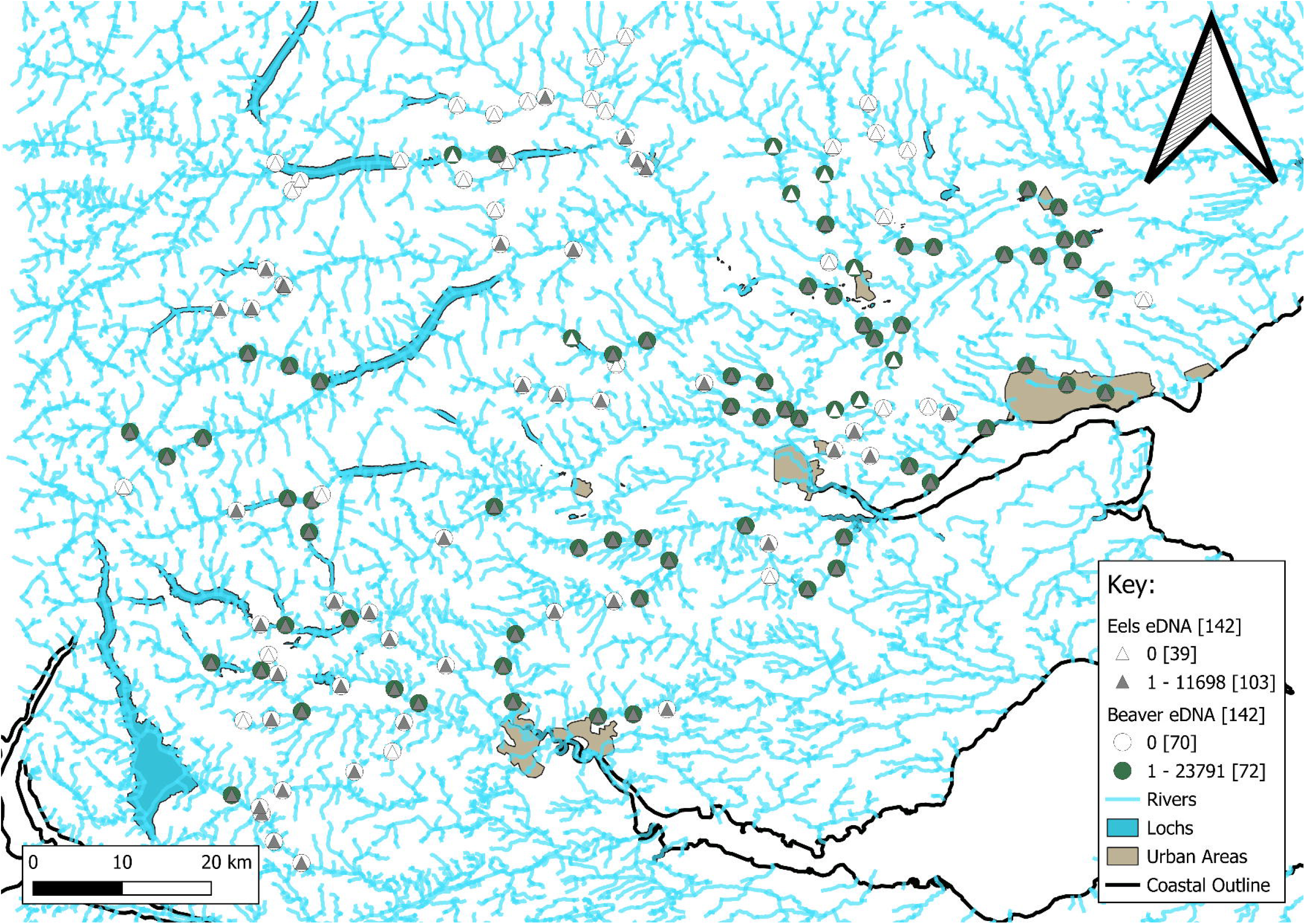

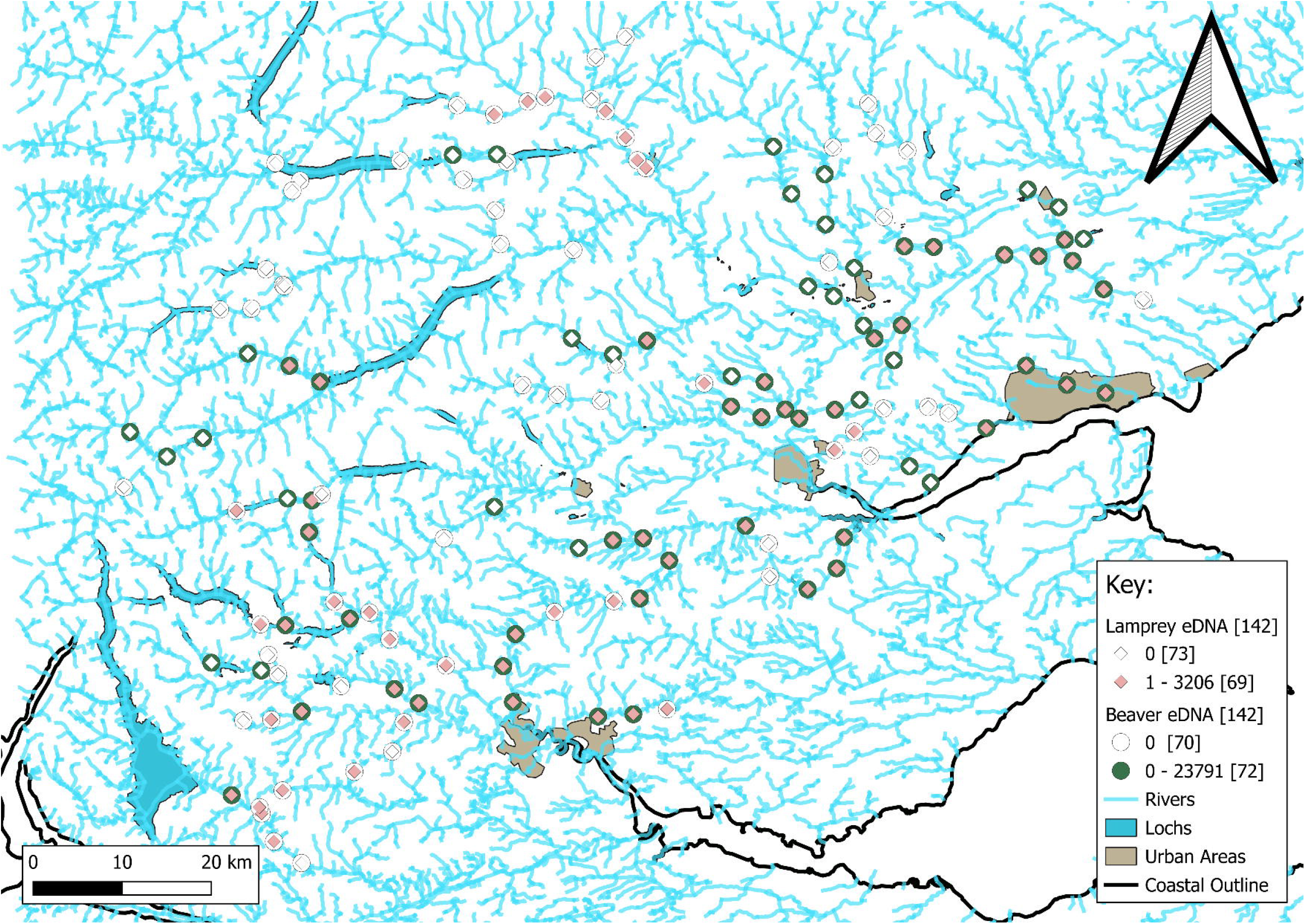

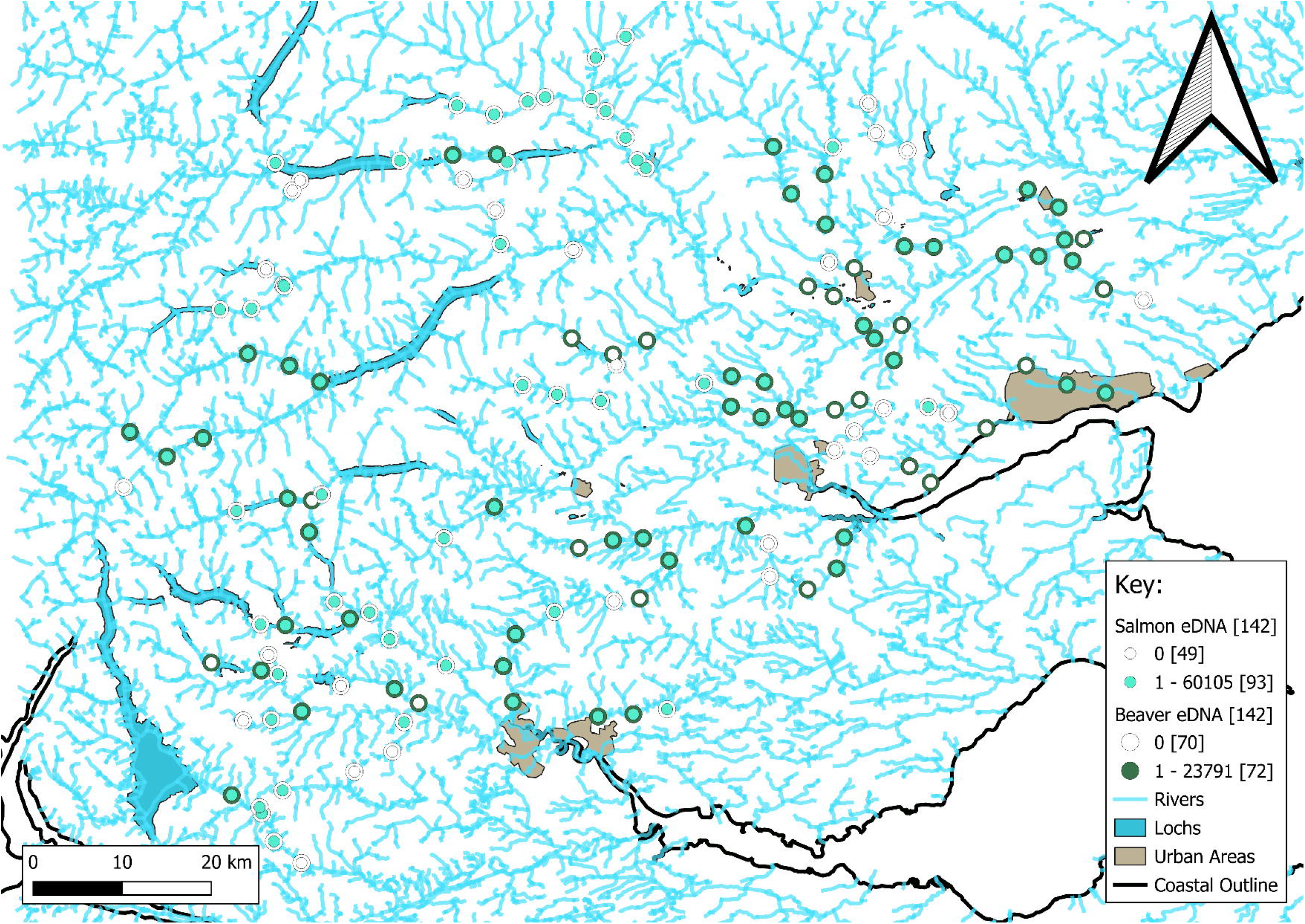

## References

Bates, D., Maechler, M., Bolker, B., & Walker, S. (2015). Fitting Linear Mixed-Effects Models Using lme4. Journal of statistical software, 67.

Beaver Salmonid Working Group. (2015). Final Report of the Beaver Salmonid Working Group. Prepared for The National Species Reintroduction Forum, Inverness.

Blackman, R. C., Carraro, L., Keck, F., & Altermatt, F. (2024). Measuring the state of aquatic environments using eDNA-upscaling spatial resolution of biotic indices. Philosophical transactions of the Royal Society of London. Series B, Biological sciences, 379, 20230121.

Brazier, R. E., Puttock, A., Graham, H. A., Auster, R. E., Davies, K. H., & Brown, C. M. L. (2021). Beaver: Nature’s ecosystem engineers. WIREs. Water, 8, e1494.

Broadhurst, H. A., Gregory, L. M., Bleakley, E. K., Perkins, J. C., Lavin, J. V., Bolton, P., … McDevitt, A. D. (2021). Mapping differences in mammalian distributions and diversity using environmental DNA from rivers. The Science of the total environment, 801, 149724.

Bull, C., Perfect, C., & Watt, J. (2016). Site condition monitoring of lamprey in the Endrick Water SSSI and SAC 2012. Scottish Natural Heritage Commissioned Report No 911.

Bylak, A., Kukuła, K., & Mitka, J. (2014). Beaver impact on stream fish life histories: the role of landscape and local attributes. Canadian journal of fisheries and aquatic sciences. Journal canadien des sciences halieutiques et aquatiques, 71, 1603–1615.

Campbell-Palmer, R., Puttock, A., Needham, R. N., Wilson, K., Graham, H., & Brazier, R. E. (2021). Survey of the Tayside Area Beaver Population 2020-2021. NatureScot Research Report 1274.

Campbell, R. D., Harrington, A., Ross, A., & Harrington, L. (2012). Distribution, population assessment and activities of beavers in Tayside. Scottish Natural Heritage Commissioned Report 540.

Cutting, K. A., Ferguson, J. M., Anderson, M. L., Cook, K., Davis, S. C., & Levine, R. (2018). Linking beaver dam affected flow dynamics to upstream passage of Arctic grayling. Ecology and evolution, 8, 12905–12917.

Darwall, W. R. T., & Noble, R. A. (2023). Salmo salar (Great Britain subpopulation). The IUCN Red List of Threatened Species, e.T213546282A213546288.

Davies, H. N., Rameshwaran, P., & Bell, V. A. (2022). Gridded (1km) physical river characteristics for the UK v2.

Di Muri, C., Handley, L. L., Bean, C. W., Li, J., Peirson, G., Sellers, G. S., … Hänfling, B. (2020). Read counts from environmental DNA (eDNA) metabarcoding reflect fish abundance and biomass in drained ponds. Metabarcoding and metagenomics, 4, e56959.

Elmeros, M., Madsen, A. B., & Berthelsen, J. P. (2003). Monitoring of reintroduced beavers (Castor fiber) in Denmark. Lutra, 46, 153–162.

Enbody, E. D., Pettersson, M. E., Sprehn, C. G., Palm, S., Wickström, H., & Andersson, L. (2021). Ecological adaptation in European eels is based on phenotypic plasticity. Proceedings of the National Academy of Sciences of the United States of America, 118, e2022620118.

Eurogeographics. (2023). EuroDEM (digital elevation model).

Falkegård, M., Lennox, R. J., Thorstad, E. B., Einum, S., Fiske, P., Garmo, Ø. A., … Forseth, T. (2023). Predation of Atlantic salmon across ontogenetic stages and impacts on populations. Canadian journal of fisheries and aquatic sciences.

Ferreira, A. F., Quintella, B. R., Maia, C., Mateus, C. S., Alexandre, C. M., Capinha, C., & Almeida, P. R. (2013). Influence of macrohabitat preferences on the distribution of European brook and river lampreys: Implications for conservation and management. Biological conservation, 159, 175–186.

Feunteun, E. (2002). Management and restoration of European eel population (Anguilla anguilla): An impossible bargain. Ecological engineering, 18, 575–591.

Fritz, S. F., & Gangloff, M. M. (2022). The effects of beaver impoundments on montane stream fish communities. Aquatic conservation: marine and freshwater ecosystems, 32, 1618–1633.

Gardiner, R., & Egglishaw, H. (1986). A map of the distribution in Scottish rivers of the Atlantic Salmon Salmo salar L.

Griffiths, N. P., Bolland, J. D., Wright, R. M., Murphy, L. A., Donnelly, R. K., Watson, H. V., & Hänfling, B. (2020). Environmental DNA metabarcoding provides enhanced detection of the European eel Anguilla anguilla and fish community structure in pumped river catchments. Journal of fish biology, 97, 1375–1384.

Griffiths, N. P., Wright, R. M., Hänfling, B., Bolland, J. D., Drakou, K., Sellers’, G. S., … Vasquez, M. I. (2023). Integrating environmental DNA monitoring to inform eel (Anguilla anguilla) status in freshwaters at their easternmost range—A case study in Cyprus. Ecology.

Hänfling, B., Lawson Handley, L., Read, D. S., Hahn, C., Li, J., Nichols, P., … Winfield, I. J. (2016). Environmental DNA metabarcoding of lake fish communities reflects long-term data from established survey methods. Molecular ecology, 25, 3101–3119.

Harper, L. R., Lawson Handley, L., Hahn, C., Boonham, N., Rees, H. C., Gough, K. C., … Hänfling, B. (2018). Needle in a haystack? A comparison of eDNA metabarcoding and targeted qPCR for detection of the great crested newt (Triturus cristatus). Ecology and evolution, 8, 6330–6341.

Harwood, A. J. P., Perrow, M. R., Sayer, C. D., Piper, A. T., Berridge, R. J., Patmore, I. R., … Cooper, G. (2022). Catchment-scale distribution, abundance, habitat use, and movements of European eel (Anguilla anguilla L.) in a small UK river: Implications for conservation management. Aquatic conservation: marine and freshwater ecosystems, 32, 797–816.

Hollis, D., McCarthy, M., Kendon, M., Legg, T., & Simpson, I. (2019). HadUK-Grid—A new UK dataset of gridded climate observations. Geoscience data journal, 6, 151–159.

Johnson, C. C., Breward, N., Ander, E. L., & Ault, L. (2005). G-BASE: baseline geochemical mapping of Great Britain and Northern Ireland. Geochemistry: Exploration, Environment, Analysis, 5, 347–357.

Kelly, R. P., Port, J. A., Yamahara, K. M., & Crowder, L. B. (2014). Using environmental DNA to census marine fishes in a large mesocosm. PloS one, 9, e86175.

Kemp, P. S., Worthington, T. A., Langford, T. E. L., Tree, A. R. J., & Gaywood, M. J. (2012). Qualitative and quantitative effects of reintroduced beavers on stream fish. Fish and fisheries, 13, 158–181.

Kemp, P. S., & O’hanley, J. R. (2010). Procedures for evaluating and prioritising the removal of fish passage barriers: a synthesis: EVALUATION OF FISH PASSAGE BARRIERS. Fisheries management and ecology, 17, 297–322.

Kitson, J. J. N., Hahn, C., Sands, R. J., Straw, N. A., Evans, D. M., & Lunt, D. H. (2019). Detecting host–parasitoid interactions in an invasive Lepidopteran using nested tagging DNA metabarcoding. Molecular ecology, 28, 471–483.

Laramie, M. B., Pilliod, D. S., & Goldberg, C. S. (2015). Characterizing the distribution of an endangered salmonid using environmental DNA analysis. Biological conservation, 183, 29–37.

Law, A., Levanoni, O., Foster, G., Ecke, F., & Willby, N. J. (2019a). Are beavers a solution to the freshwater biodiversity crisis? Diversity & distributions, 25, 1763–1772.

Law, A., Baker, A., Sayer, C., Foster, G., Gunn, I. D. M., Taylor, P., … Willby, N. J. (2019b). The effectiveness of aquatic plants as surrogates for wider biodiversity in standing fresh waters. Freshwater biology, 64, 1664–1675.

Lokteff, R. L., Roper, B. B., & Wheaton, J. M. (2013). Do Beaver Dams Impede the Movement of Trout? Transactions of the American Fisheries Society, 142, 1114–1125.

Lüdecke, M. D. (2024). sjPlot: Data Visualization for Statistics in Social Science. R package version 2.8.16 2024.

Malcolm, I. A., Millidine, K. J., Glover, R. S., Jackson, F. L., Millar, C. P., & Fryer, R. J. (2019a). Development of a large-scale juvenile density model to inform the assessment and management of Atlantic salmon (Salmo salar) populations in Scotland. Ecological indicators, 96, 303–316.

Malcolm, I. A., Jackson, F. L., Millidine, K. J., Bacon, P. J., McCartney, A. G., & Fryer, R. J. (2023). The National Electrofishing Programme for Scotland (NEPS) 2021. Scottish Marine and Freshwater Science Reports, 14.

Malcolm, I. A., Millidine, K. J., Jackson, F. L., Glover, R. S., & and Fryer, R. J. (2019b). Assessing the status of Atlantic salmon (Salmo salar) from juvenile electrofishing data collected under the National Electrofishing Programme for Scotland (NEPS). Scottish Marine and Freshwater Science, 10.

Malison, R. L., & Halley, D. J. (2020). Ecology and movement of juvenile salmonids in beaver-influenced and beaver-free tributaries in the Trøndelag province of Norway. Ecology of freshwater fish, 29, 623–639.

Marine Directorate. (2008). Salmon rivers in Scotland, 2008.

Marshall Wolf, J., Clancy, N. G., & Rosenthal, L. R. (2024). Bull Trout Passage at Beaver Dams in Two Montana Streams. Northwest Science, 97, 68–77.

Moore, R. V., Morris, D. G., & Flavin, R. W. (1994). Sub-set of UK digital 1: 50,000 scale river centre-line network. NERC, Institute of Hydrology, Wallingford.

Morton, R. D., Marston, C. G., O’Neil, A. W., & Rowland, C. S. (2024). Land Cover Map 2023 (10m classified pixels, GB).

Naiman, R. J., Melillo, J. M., & Hobbie, J. E. (1986). Ecosystem alteration of boreal forest streams by beaver (Castor canadensis). Ecology, 67, 1254–1269.

Needham, R. J., Gaywood, M., Tree, A., Sotherton, N., Roberts, D., Bean, C. W., & Kemp, P. S. (2021). The response of a brown trout (Salmo trutta) population to reintroduced Eurasian beaver (Castor fiber) habitat modification. Canadian journal of fisheries and aquatic sciences. Journal canadien des sciences halieutiques et aquatiques, 78, 1650– 1660.

Needham, R. J., Zabel, R. W., Roberts, D., & Kemp, P. S. (2025). The impact of reintroduced Eurasian beaver (Castor fiber) dams on the upstream movement of brown trout (Salmo trutta) in upland areas of Great Britain. PloS one, 20, e0313648.

Nunn, A. D., Ainsworth, R. F., Walton, S., Bean, C. W., Hatton-Ellis, T. W., Brown, A., … Noble, R. A. A. (2023). Extinction risks and threats facing the freshwater fishes of Britain. Aquatic conservation: marine and freshwater ecosystems, 33, 1460–1476.

Pander, J., Otterbein, J., Nagel, C., & Geist, J. (2025). Nature-based fish habitat enrichment of non-damming beaver structures positively affects fish species richness and density. Ecological engineering, 212, 107516.

Piper, A. T., Wright, R. M., Walker, A. M., & Kemp, P. S. (2013). tEscapement, route choice, barrier passage and entrainment of seaward migrating European eel, Anguilla anguilla, within a highly regulated lowland river. Ecological engineering, 57, 88–96.

Pont, D. (2024). Predicting downstream transport distance of fish eDNA in lotic environments. Molecular ecology resources, 24, e13934.

Pont, D., Rocle, M., Valentini, A., Civade, R., Jean, P., Maire, A., … Dejean, T. (2018). Environmental DNA reveals quantitative patterns of fish biodiversity in large rivers despite its downstream transportation. Scientific reports, 8.

QGIS Development Team. (2025). QGIS geographic information system.

Quail, M. A., Swerdlow, H., & Turner, D. J. (2009). Improved protocols for the illumina genome analyzer sequencing system. Current protocols in human genetics, Chapter 18, Unit 18.2.

R Core Team. (2025). R: A Language and Environment for Statistical Computing. R Foundation for Statistical Computing, Vienna, Austria.

Riaz, T., Shehzad, W., Viari, A., Pompanon, F., Taberlet, P., & Coissac, E. (2011). ecoPrimers: inference of new DNA barcode markers from whole genome sequence analysis. Nucleic acids research, 39, e145.

Sellers, G. S., Di Muri, C., Gómez, A., & Hänfling, B. (2018). Mu-DNA: a modular universal DNA extraction method adaptable for a wide range of sample types. Metabarcoding and metagenomics, 2, e24556.

SEPA-WMS. (2018). Obstacles to Fish Passage.

Solomon, D., & Beach, M. H. (2004). Fish pass design for eel and elver (Anguilla anguilla). R&D Technical Report W2-070/TR1. Bristol: Environment Agency.

Soulsby, C., Youngson, A. F., Moir, H. J., & Malcolm, I. A. (2001). Fine sediment influence on salmonid spawning habitat in a lowland agricultural stream: a preliminary assessment. The Science of the total environment, 265, 295–307.

Stringer, A. P., & Gaywood, M. J. (2016). The impacts of beavers Castor spp. on biodiversity and the ecological basis for their reintroduction to Scotland, UK. Mammal review, 46, 270–283.

Stringer, A. P., Blake, D., Genney, D. R., & Gaywood, M. J. (2018). A geospatial analysis of ecosystem engineer activity and its use during species reintroduction. European journal of wildlife research, 64.

Wickham, H. (2016). ggplot2: Elegant Graphics for Data Analysis. Springer-Verlag New York 2016.

Worldwide Fund for Nature. (2024). 2024 living planet report: A System in Peril. Gland: WWF.

