## Supplementary material for "eDNA sampling reveals no negative effects of beaver recolonisation on the catchment-scale distribution of migratory fish": Word document containing all supporting information for the publication

**Supplementary 1: Detailed metabarcoding workflow**

**Sample collection**

Across our sites 426 2L surface water samples were collected and filtered following the methodology described in Griffiths *et al.,* (2023). Sampling was carried out from the 06/07/2023 – 18/08/2023 and at each site three 2-Litre water samples were taken using sterile Gosselin HDPE plastic bottles (Fisher Scientific UK Ltd., Loughborough, UK), sampling equipment was sterilised between sites using 10% bleach and a maximum of 12 sites were visited each day. A field blank consisting of 2-L purified water was used each day (n=19), this was taken into the field and handled alongside eDNA samples to monitor for contamination. Following collection samples were placed on ice packs within bleach sterilised coolboxes before being taken back to an eDNA filtration facility at the University of Stirling.

**Filtration**

Water samples were vacuum filtered within 24 hours of collection following the methodology described in Griffiths *et al*., (2023). Briefly, 2-L samples were vacuum filtered through sterile 0.45 μm mixed cellulose nitrate membrane filters with pads (47 mm diameter; Whatman, GE Healthcare) using Pall filtration units. Two filters were used for each sample to minimise clogging, and up to 30 minutes per filter was allowed for 1-L of water to pass through. Filters were removed from pads using sterile tweezers, rolled, and placed back-to-back in sterile 5 ml Axygen screw-cap tubes (Fisher Scientific UK Ltd., Loughborough, UK) before being stored at -20 °C. Up to nine samples were filtered each run and blanks (n=19) were filtered during the last round of each day. Between runs, attachments were submerged in 10% v/v chlorine-based bleach solution (Fisher Scientific UK Ltd., Loughborough, UK) for 10 min, soaked in 5% v/v MicroSol detergent (Scientific Laboratory Supplies, Newhaven, UK) for a further 5 min, before being rinsed carefully with purified water to remove any remaining detergent. Throughout filtration, all benchtops and equipment within the filtration lab were sterilised using 10% bleach solution and cleaned using 70% v/v ethanol (Fisher Scientific UK Ltd., Loughborough, UK) (Griffiths *et al.*, 2023)**.**

**DNA extractions**

DNA extractions were carried out following the mu-DNA lysis water protocol described in Sellers *et al*., (2018) but with the following modifications: instead of garnet beads, the lysis stage used an incubation based method where 34 μl of 20 mg/mL proteinase K (VWR, Leicestershire, UK) was added to 750 μl of lysis solution and 250 μl of water lysis additive. The filter was then pressed to the bottom of the tube using the pipette tip until it was submerged below the liquid before tubes were incubated at 55°C overnight. The ethanol wash step was repeated twice to minimise the chances of inhibition before the rest of the protocol was followed as normal (Sellers *et al.*, 2018). An extraction blank (n=20), consisting only of extraction buffers, was extracted alongside samples to monitor for contamination. Following extraction, the eluted DNA extracts (100 μL) were visualised using the QIAxpert (Qiagen, Hilden, Germany) to confirm DNA was successfully isolated, then stored at −20°C.

**Metabarcoding**

Library preparation and sequencing followed a modified version of the established 12S metabarcoding protocols described in Griffiths *et al*., (2023). Briefly, nested metabarcoding was carried out following a two-step PCR approach in which both PCRs used multiplex identification tags to enable sample identification following the protocols described in Kitson *et al*., (2019). A positive control (quantified at 0.05 ng/μl) from the non-native cichlid (*Astatotilapia calliptera*) and a negative control of molecular grade water were used for each library. The first PCR was carried out in triplicate (3x PCR replicates) to amplify a 106bp fragment using published 12S ribosomal RNA primers 12S-V5-F (5’ -ACTGGGATTAGATACCCC-3’) and 12S-V5-R (5’ -TAGAACAGGCTCCTCTAG-3’ ) (Kelly *et al.*, 2014; Riaz *et al.*, 2011). These primers have been validated both *in vitro, in silico* and *in situ* for UK vertebrates (Hänfling *et al.*, 2016), confirming that for the purpose of the current study, all UK migratory fish species can be detected with the exceptions of distinctions between *Lampetra planeri* / *Lampetra fluviatilis* which will be assigned to the genus *Lampetra*.

Following the first PCR, samples were visualised on a zero-agarose gel (ZAG) DNA Analyzer System (Agilent Technologies) using the ZAG 135 dsDNA Kit (1-1500bp) (Agilent Technologies). Samples were then normalised within PCR1 plates based on band strength as follows: 5µl - very bright band, 10 µl - bright band, 15 µl - weak band, 20 µl - no band. Subsequently, to remove non-specific amplification, libraries underwent a single size selection clean-up (Quail *et al.*, 2009). The first PCR underwent a single size selection protocol using the ProNex® Size-Selective Purification beads at a ratio of 1.5 x 100µl to remove amplification below the target band. A second PCR was carried out to bind pre adapters, indexes, and Illumina adapters to each sub library. This was carried out in duplicate (2 x PCR replicates) following the adapted thermocycling profile described in Griffiths et al., (2023). Duplicate PCR products were then pooled and visualised following the conditions described previously. A double size selection clean-up (Quail *et al.*, 2009) was then carried out to remove primer dimer and nonspecific amplification at the ratios of 1.2 x and 0.4 x magnetic beads to 50μl of PCR product.

Cleaned sub-libraries were quantified using the Qubit 3.0 fluorometer high-sensitivity (HS) dsDNA assay (Invitrogen), each library was diluted to 5ng/μl and pooled proportionally according to sample number. The library was then diluted to 4nM before being quantified via qPCR using the Collibri™ Library Quantification Kit for Illumina (Roche, Hertfordshire, UK). Once the desired quantification was confirmed, the final library was denatured and sequenced at 13pM with 10% PhiX Control on an Illumina MiSeq using a MiSeq Reagent Kit v3 (600 cycle) (Illumina Inc., San Diego, CA, USA) (Griffiths *et al.*, 2023).

**Methods References:**

Griffiths, N. P., Wright, R. M., Hänfling, B., Bolland, J. D., Drakou, K., Sellers’, G. S., … Vasquez, M. I. (2023). Integrating environmental DNA monitoring to inform eel (*Anguilla anguilla*) status in freshwaters at their easternmost range—A case study in Cyprus. *Ecology*.

Kitson, J. J. N., Hahn, C., Sands, R. J., Straw, N. A., Evans, D. M., & Lunt, D. H. (2019). Detecting host–parasitoid interactions in an invasive Lepidopteran using nested tagging DNA metabarcoding. *Molecular ecology*, *28*, 471–483.

Quail, M. A., Swerdlow, H., & Turner, D. J. (2009). Improved protocols for the illumina genome analyzer sequencing system. *Current protocols in human genetics*, *Chapter 18*, Unit 18.2.

Sellers, G. S., Di Muri, C., Gómez, A., & Hänfling, B. (2018). Mu-DNA: a modular universal DNA extraction method adaptable for a wide range of sample types. *Metabarcoding and metagenomics*, *2*, e24556.

**Supplementary 2: Full list of model covariates alongside their appropriate spatial datasets**

| **Covariate (Abbreviation)** | **Dataset and appropriate sources** | **Unit** | **Description** |
| --- | --- | --- | --- |
| Impassable barriers (IPB) | SEPA obstacles to fish passage (SEPA-WMS, 2018) |  | Number of impassable barriers downstream |
| Passable barriers - fish pass (passable barriers - FP) | SEPA obstacles to fish passage (SEPA-WMS, 2018) |  | Number of passable barriers with fish passes downstream |
| Passable barriers - no fish pass (passable barriers - NFP) | SEPA obstacles to fish passage (SEPA-WMS, 2018) |  | Number of passable barriers without fish passes downstream |
| Distance from coast | Ordnance Survey Open Rivers | KM | Distance from each point to the coast measured along the river |
| Elevation | EuroDEM (Eurogeographics, 2023) | M | Height of each site above sea level |
| Slope | EuroDEM (Eurogeographics, 2023) | Degrees | The gradient at each site from horizontal |
| Upstream catchment area (upstream catchment area) | Gridded (1km) physical river characteristics for the UK v2 (Davies *et al.*, 2022) | KM2 | Catchment area upstream of a site - Upstream catchment area was taken from 1 km x 1 km resolution raster layers of gridded physical river habitat characteristics, GPS data from sampling sites that could not snap to this grid were adjusted accordingly |
| Bankfull river width | Gridded (1km) physical river characteristics for the UK v2 (Davies *et al.*, 2022) | M | Bankfull width of the river channel at each site - bankfull river width was taken from 1 km x 1 km resolution raster layers of gridded physical river habitat characteristics, GPS data from sampling sites that could not snap to this grid were adjusted accordingly |
| pH | 92 - in field  50 sites - G-Base stream layer (Johnson *et al.*, 2005) |  | pH of the water at each site - For the 50 sites where it was not possible to collect water pH in the field, the nearest appropriate point was selected from the BGS (British Geological Survey) G-Base stream layer |
| Mean atmospheric temperature 2023 | HadUK-Grid Gridded Climate Observations on a 1km grid over the UK, v1.3.0.ceda (1836-2023) (Hollis *et al.*, 2019) | (℃) | Mean annual atmospheric temperature in 2023 |
| Total annual rainfall 2023 | HadUK-Grid Gridded Climate Observations on a 1km grid over the UK, v1.3.0.ceda (1836-2023) (Hollis *et al.*, 2019) | (mm) | Total annual precipitation in 2023 |
| Northern Pike (*Esox lucius*) | 2023 eDNA data | 1 (Presence) and -1 (Absence) | Presence/absence data for Northern pike |
| Eurasian beaver | 2023 eDNA data | 1 (Presence) and -1 (Absence) | Presence/absence data for Eurasian beaver |
| Local land-use | CEH Land cover map 2023 10m raster (Morton *et al.*, 2024) | % | % cover of different land-use categories from 500 m circular buffer x 25 m either side of the river |
| Upstream land-use | CEH Land cover map 2023 10m raster (Morton *et al.*, 2024) | % | % cover of different land-use categories from 1 km upstream buffer x 25 m of the river |
| Catchment | EuroDEM (Eurogeographics, 2023) |  | River catchment which each site belongs to - River catchments were generated through a hydrological delineation of watersheds using the “channel network and drainage basins” plugin in the SAGA toolbox (Conrad et al., 2015) and the EuroDEM digital elevation model (Eurogeographics, 2023). For the smaller catchments, such as Pitroddie and Huntly, where the DEM did not align well, the appropriate river catchment was corrected. |

**References:**

Davies, H. N., Rameshwaran, P., & Bell, V. A. (2022). Gridded (1km) physical river characteristics for the UK v2.

Eurogeographics. (2023). EuroDEM (digital elevation model).

Hollis, D., McCarthy, M., Kendon, M., Legg, T., & Simpson, I. (2019). HadUK‐Grid—A new UK dataset of gridded climate observations. *Geoscience data journal*, *6*, 151–159.

Johnson, C. C., Breward, N., Ander, E. L., & Ault, L. (2005). G-BASE: baseline geochemical mapping of Great Britain and Northern Ireland. *Geochemistry: Exploration, Environment, Analysis*, *5*, 347–357.

Morton, R. D., Marston, C. G., O’Neil, A. W., & Rowland, C. S. (2024). Land Cover Map 2023 (10m classified pixels, GB).

SEPA-WMS. (2018). Obstacles to Fish Passage.

**Supplementary 3: Final model covariates alongside descriptive statistics and which of the parsimonious final models each covariate was retained**

| Covariate | Mean (standard deviation) | Range | Final model  (E = Eels, S = Salmon, L = Lamprey) |
| --- | --- | --- | --- |
| Impassable barriers | 0.169 (± 0.412) | 0 - 2 | E, L |
| Passable barriers - FP | 0.704 (± 1.016) | 0 - 4 | E |
| Passable barriers - NFP | 1.627 (± 1.486) | 0 - 5 | E |
| Total Rainfall (mm) | 1412 (± 385.8) | 875.1 - 2833 | E, S |
| Elevation (M) | 123.3 (± 88.03) | 8 - 416 | E, L |
| Distance from coast (KM) | 63.01 (± 32.6) | 1.53 - 142.8 | E, L |
| Upstream catchment area (KM) | 128.9 (± 234.3) | 1 - 165 | E, S, L |
| Broadleaved woodland (%) - Upstream land use | 28.03 (± 24.4) | 0 - 91.6 | L |
| Coniferous woodland (%) - Upstream land use | 5.21 (± 11.4) | 0 - 68.6 | S, L |
| Agricultural (%) - Upstream land use | 46.32 (± 28) | 0.4 - 100 | S, L |

**Supplementary 4: Model AIC values throughout the stepwise selection process**

| **Model and dropped term** | **AIC** |
| --- | --- |
| **Salmon** |  |
| Full Model - without random effect | 117.18 |
| Full Model - with random effect | 119.2 |
| pH | 115.18 |
| Elevation | 113.19 |
| Wetland | 111.21 |
| Passable barriers with fish pass | 109.23 |
| Broadleaved woodland | 107.27 |
| Distance from coast | 105.59 |
| Beavers | 104.08 |
| Urban areas | 102.86 |
| Passable barriers without fish pass | 102.11 |
| **Lamprey** |  |
| Full model - without random effect | 142.33 |
| Full model - with random effect | 144.3 |
| Urban areas | 140.33 |
| Passable barriers - without fish pass | 138.34 |
| Pike | 136.43 |
| Beavers | 134.84 |
| Passable barriers with fish passes | 134.26 |
| pH | 133.59 |
| Total Rainfall | 133.46 |
| **Eels** |  |
| Full model - without random effect | 98.817 |
| Full model - with random effect | 100.8 |
| Agricultural | 96.86 |
| pH | 94.909 |
| Pike | 93.48 |
| Broadleaved woodland | 92.29 |
| Coniferous woodland | 91.306 |
| Wetland | 90.843 |
| Urban | 89.824 |

**Supplementary 5: Final model summaries**

| **Covariate** | **Estimate** | **Standard Error** | **Z-value** | **P-value** | **Significance** |
| --- | --- | --- | --- | --- | --- |
| **Salmon** |  |  |  |  |  |
| (Intercept) | 2.171789 | 0.400236 | 5.426273 | 5.75E-08 | ** |
| SDM.UCAmax | 1.457021 | 0.335517 | 4.342609 | 1.41E-05 | *** |
| SDM.Tot_Rain | 0.537281 | 0.338899 | 1.585371 | 0.112882 |  |
| SDM.Pike | -2.19398 | 0.797585 | -2.75077 | 0.005945 | ** |
| Agricultural | -0.51691 | 0.286673 | -1.80312 | 0.071369 |  |
| SDM.CF_Wood | -0.44879 | 0.30208 | -1.48567 | 0.137367 |  |
| **Lamprey** |  |  |  |  |  |
| (Intercept) | -0.38249 | 0.259615 | -1.47332 | 0.140666 |  |
| SDM.IPB | -0.74966 | 0.363385 | -2.06299 | 0.039114 | * |
| SDM.UCAmax | 1.360041 | 0.32331 | 4.206621 | 2.59E-05 | *** |
| SDM.Elevation | -0.73162 | 0.327255 | -2.23563 | 0.025376 | * |
| SDM.Distance | -0.71864 | 0.295674 | -2.43053 | 0.015077 | * |
| Agricultural | 1.484624 | 0.440987 | 3.366593 | 0.000761 | *** |
| SDM.BR_Wood | 1.164721 | 0.39027 | 2.984402 | 0.002841 | ** |
| SDM.CF_Wood | 0.578133 | 0.303496 | 1.904915 | 0.056791 |  |
| **Eels** |  |  |  |  |  |
| (Intercept) | 2.534827 | 0.591802 | 4.283234 | 1.84E-05 | *** |
| SDM.IPB | -1.22315 | 0.368924 | -3.31546 | 0.000915 | *** |
| SDM.PB.FP | -1.28695 | 0.415252 | -3.0992 | 0.00194 | ** |
| SDM.PB.NFP | 1.072658 | 0.597307 | 1.795823 | 0.072523 |  |
| SDM.upstream catchment areamax | 0.853678 | 0.346113 | 2.466471 | 0.013645 | * |
| SDM.Elevation | -2.33185 | 0.868292 | -2.68556 | 0.007241 | ** |
| SDM.Distance | -0.98888 | 0.620796 | -1.59292 | 0.111178 |  |
| SDM.Tot_Rain | 1.385945 | 0.469619 | 2.951211 | 0.003165 | ** |
| SDM.Beavers | 0.77093 | 0.38235 | 2.016291 | 0.04377 | * |

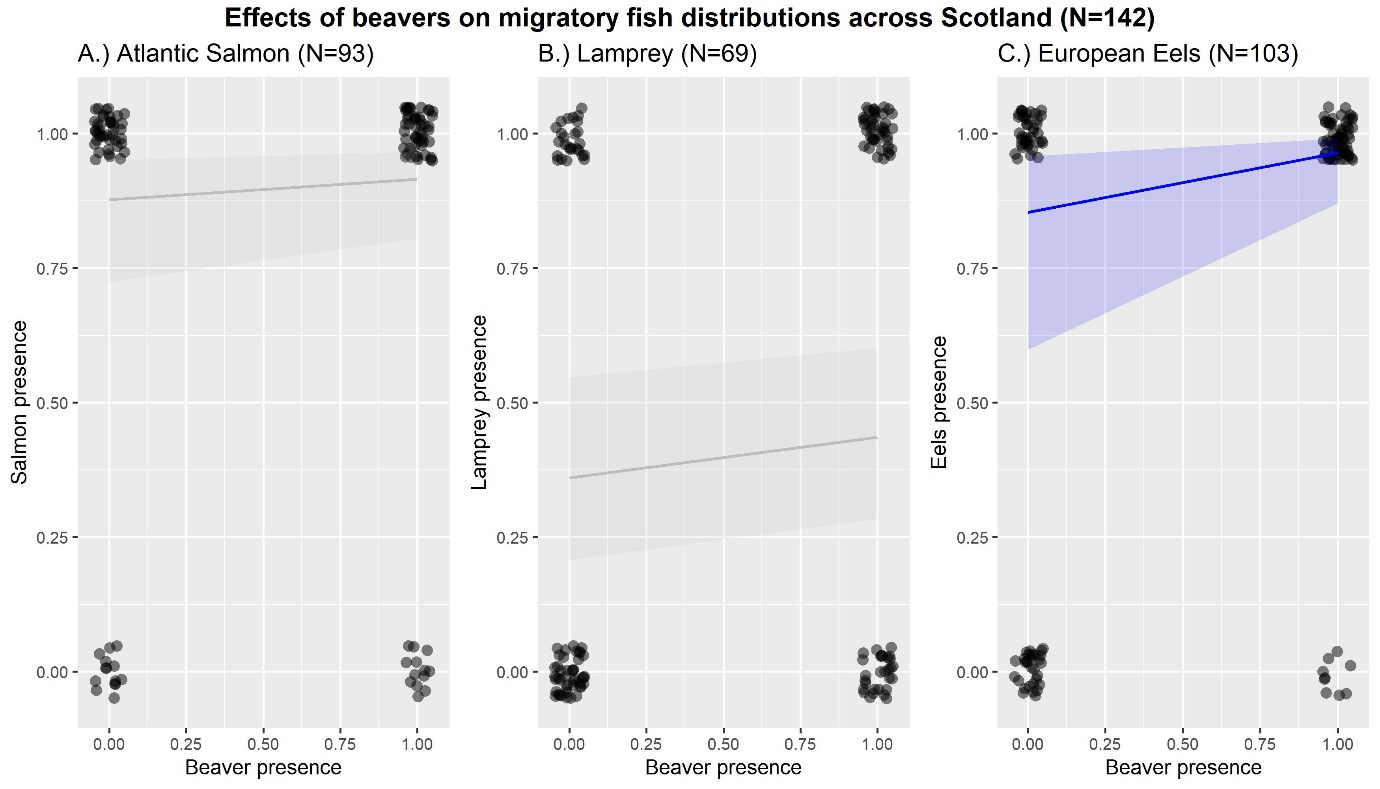

**Supplementary 6: The effects of beavers on the 3 migratory fish species before beavers are dropped from the salmon and lamprey models, the lines are coloured based on significance (P <0.05) (non-significant interactions = grey, significant interactions = blue) and the shading represents the 95% confidence interval.**

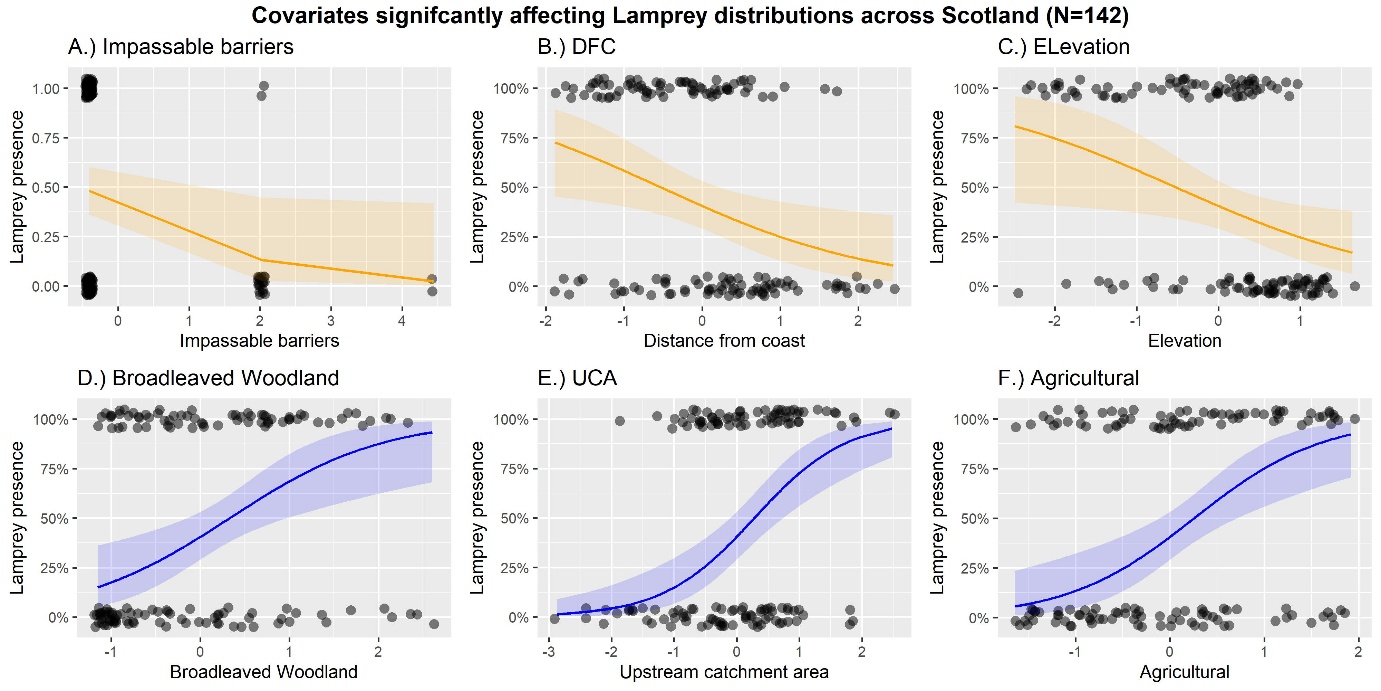

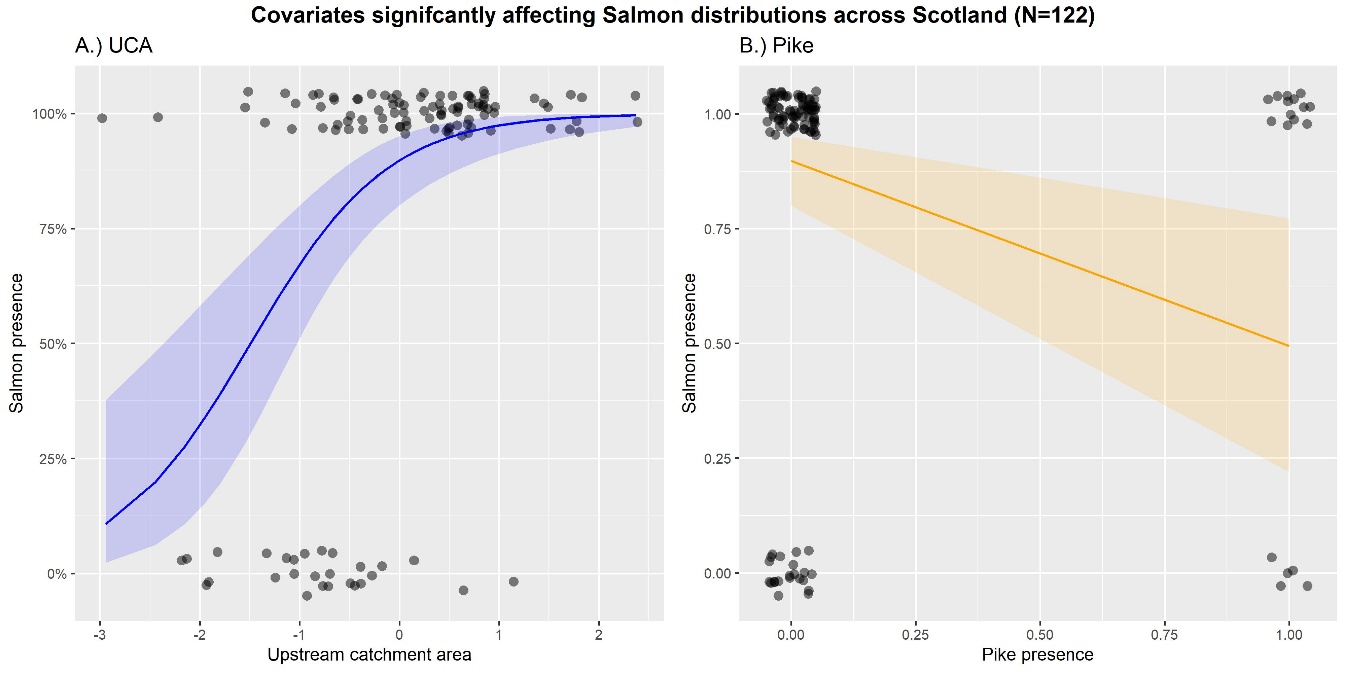
**Supplementary 7: Covariates significantly affecting salmon distribution (P <0.05) (Positive effect = blue, Negative effect = orange) and the shading represents the 95% confidence interval.**

**Supplementary 8: Covariates significantly affecting Lamprey distribution (P <0.05) (Positive effect = blue, Negative effect = orange) and the shading represents the 95% confidence interval.**

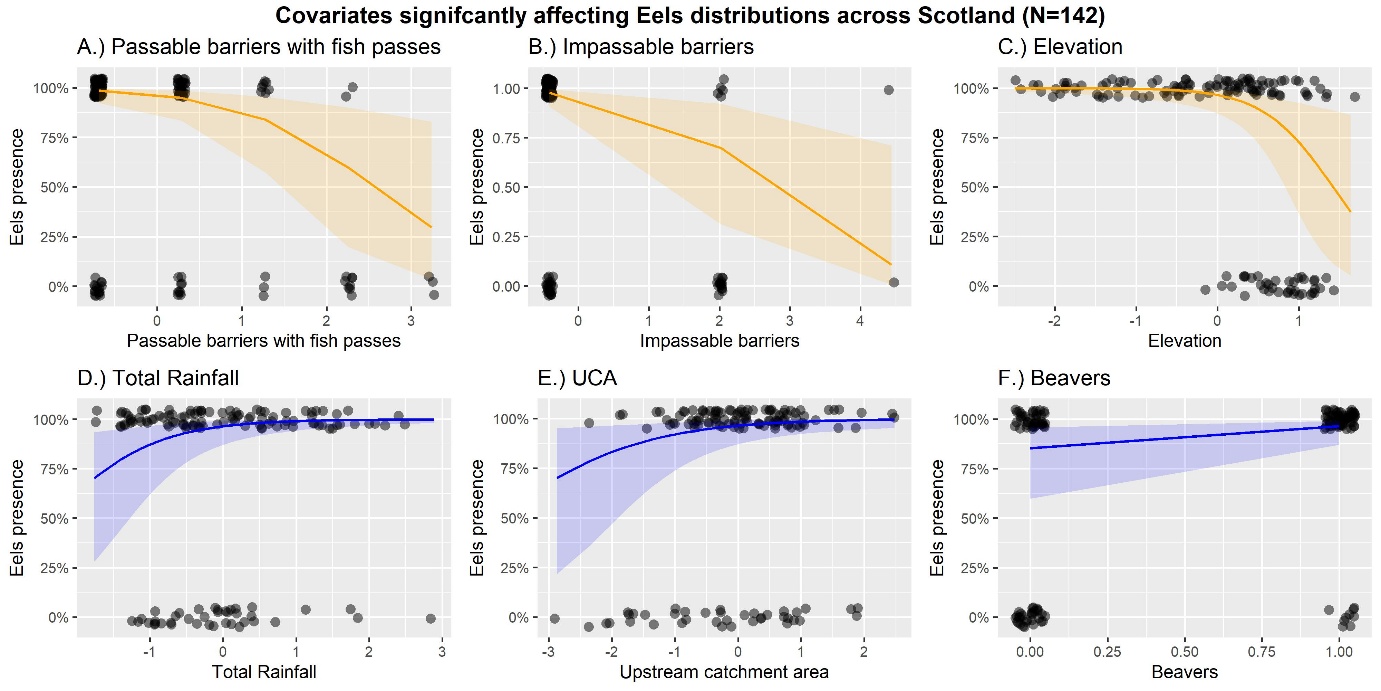
**Supplementary 9: Covariates significantly affecting eel distribution (P <0.05) (Positive effect = blue, Negative effect = orange) and the shading represents the 95% confidence interval.**

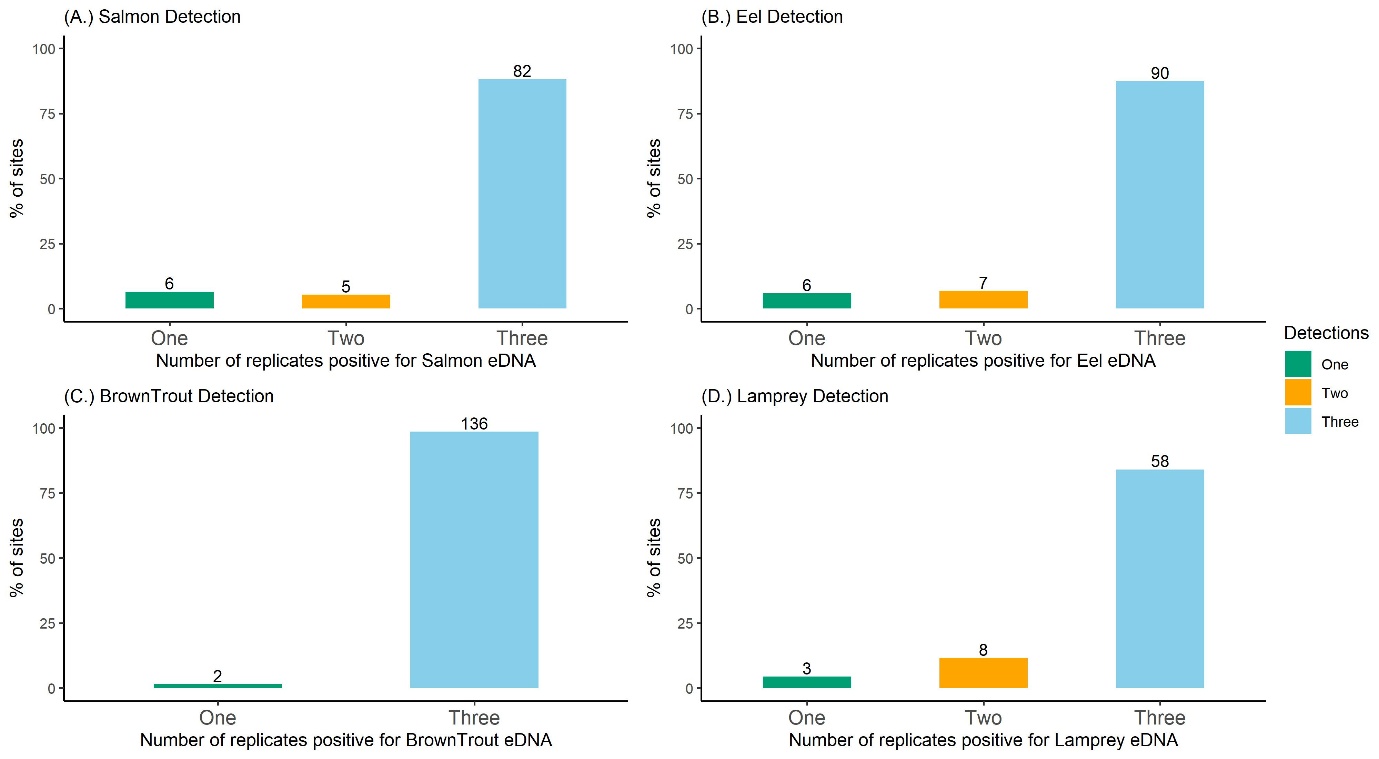

**Supplementary 10: Species detections across the three biological replicates for the 4 migratory fish species. Bars are coloured based on the number of replicates positive (one = green, two = yellow, three = blue) and the numbers above the bars represent the number of positive sites.**

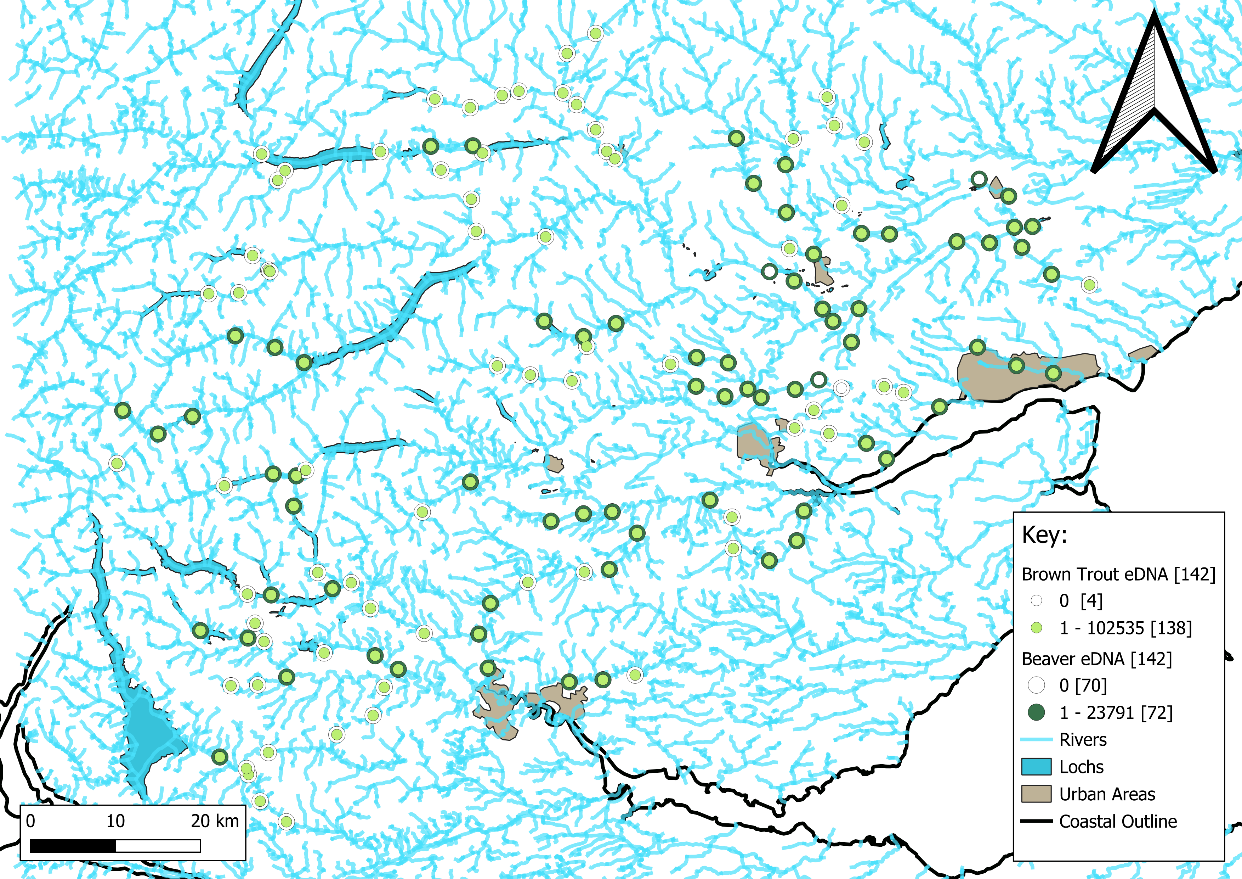

**Supplementary 11: Brown trout (n = 138) and Beaver (n = 72) positive eDNA sites across the 142 sampling locations.**

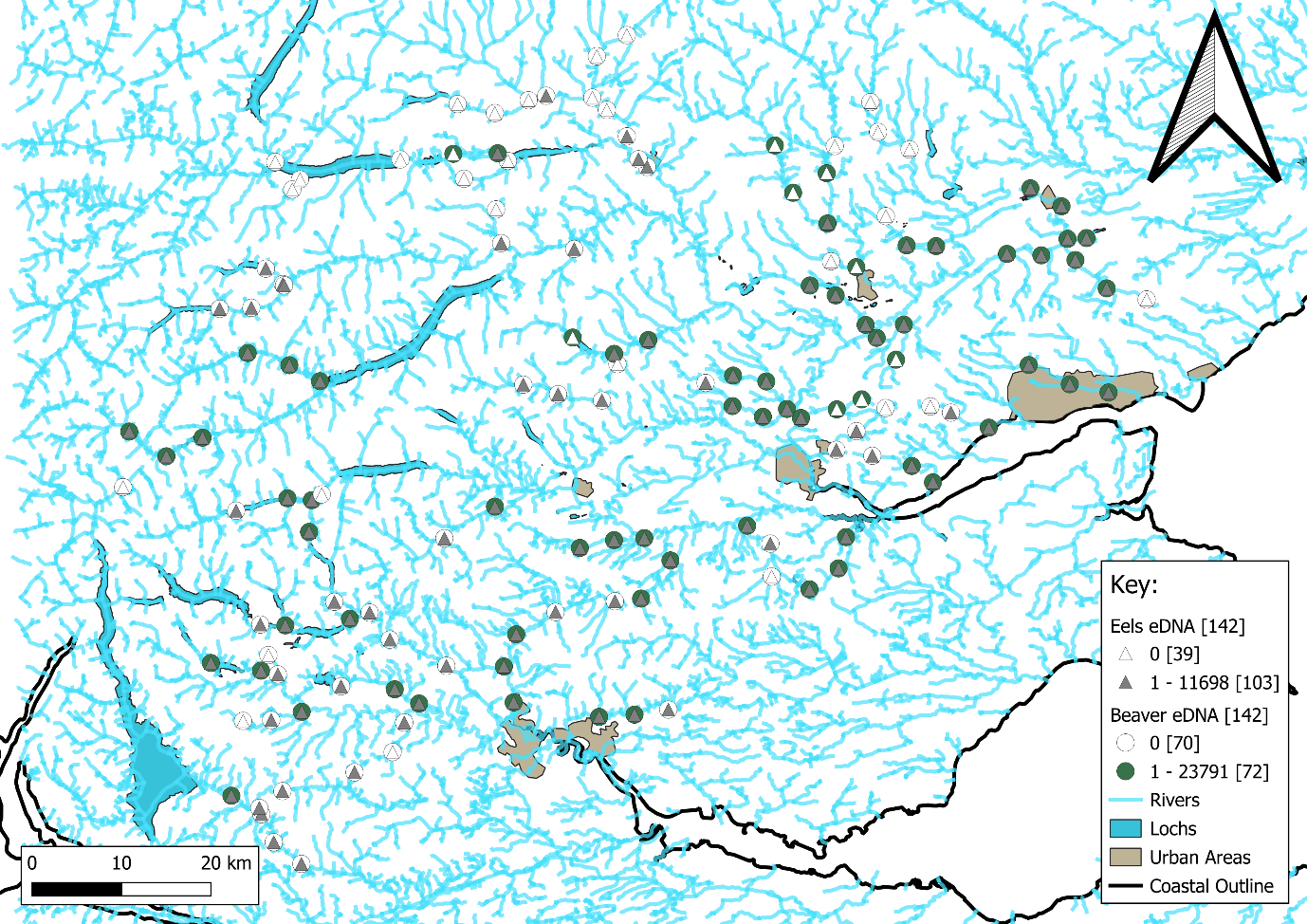
**Supplementary 12: European eel (n = 103) and Beaver (n = 72) positive eDNA sites across the 142 sampling locations.**

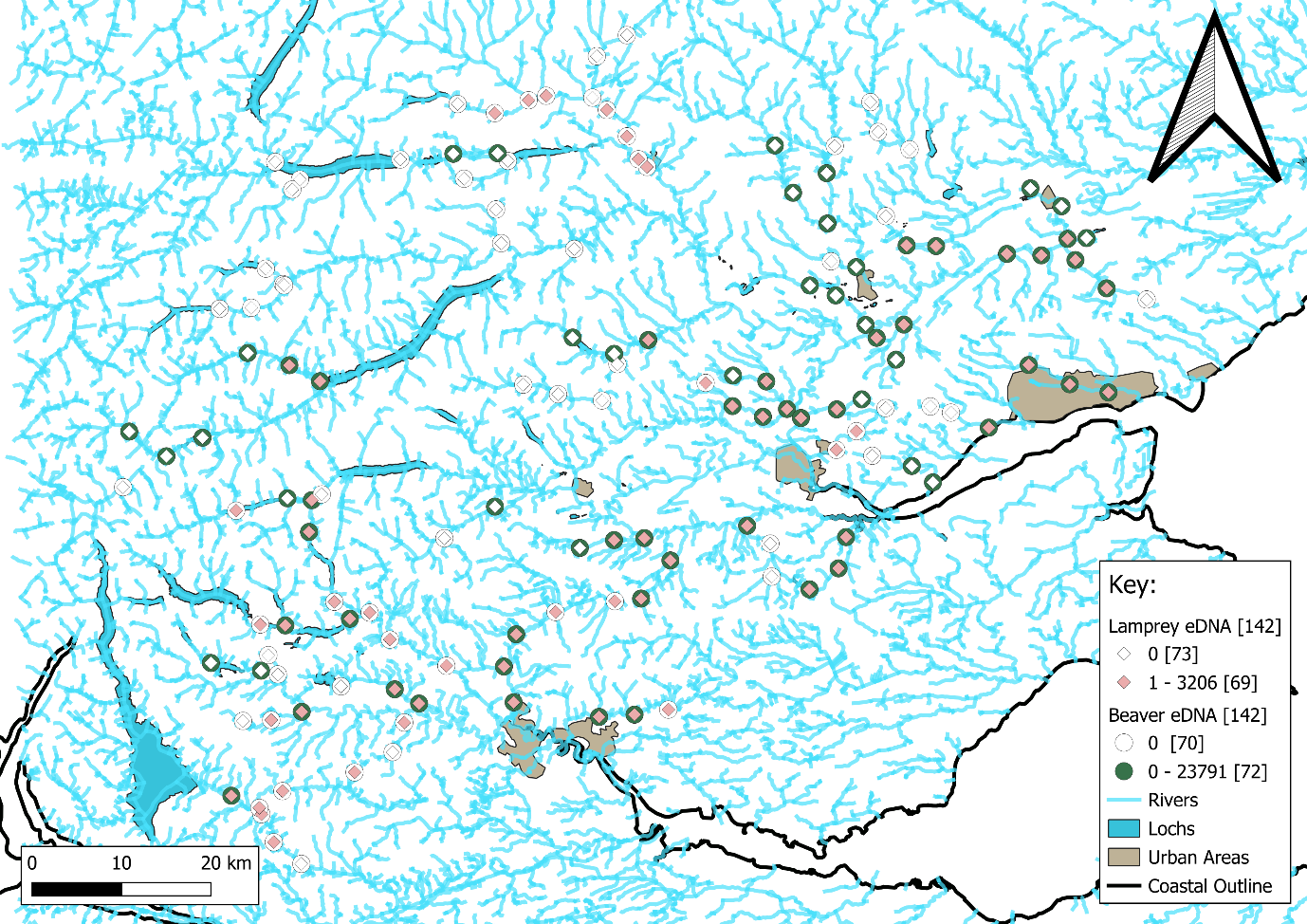
**Supplementary 13: Lamprey (n = 69) and Beaver (n = 72) positive eDNA sites across the 142 sampling locations.**

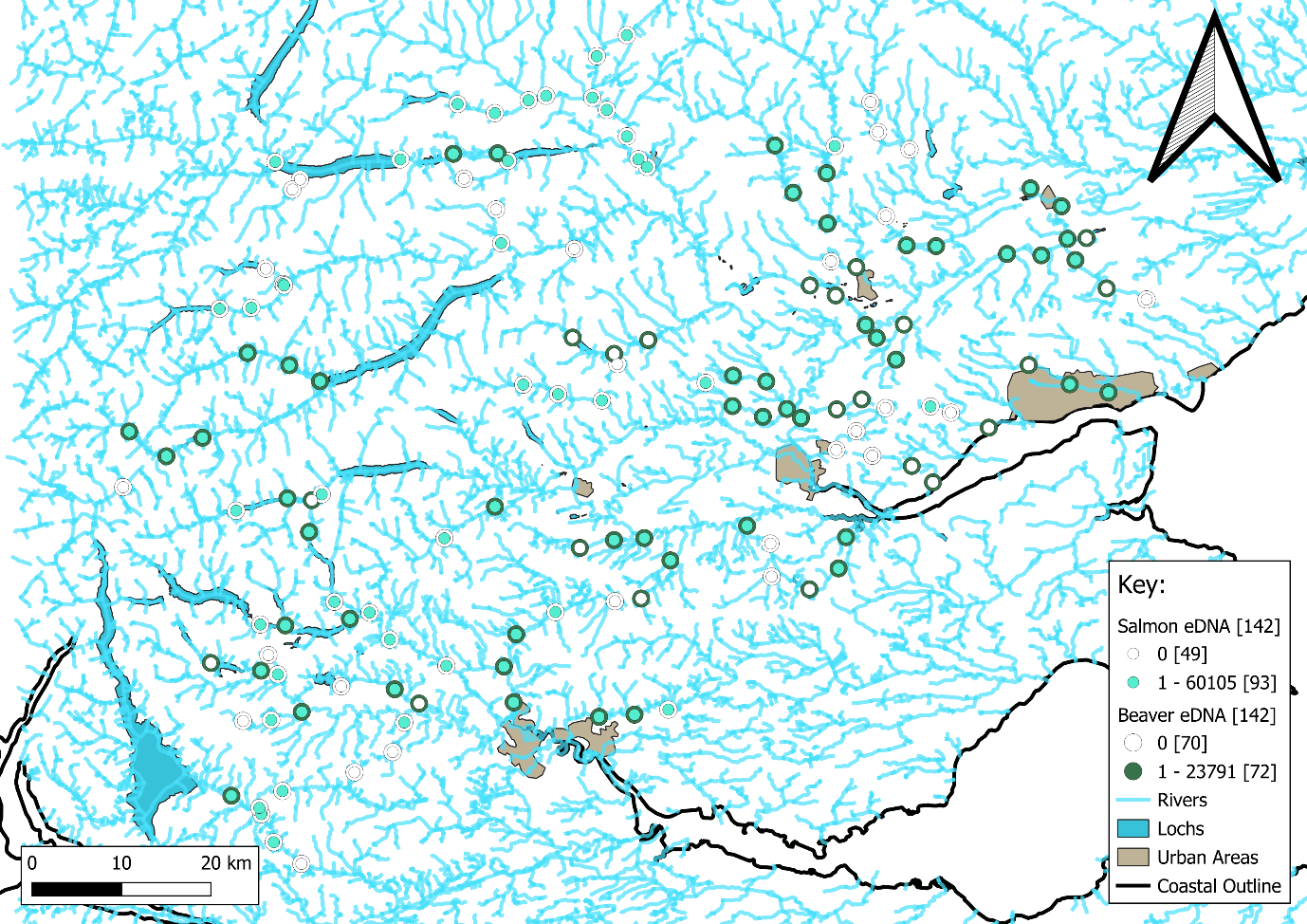
**Supplementary 14: Atlantic salmon (n = 103) and Beaver (n = 72) positive eDNA sites across the 142 sampling locations.**
